# ModelRevelator: Fast phylogenetic model estimation via deep learning

**DOI:** 10.1101/2021.12.22.473813

**Authors:** Sebastian Burgstaller-Muehlbacher, Stephen M. Crotty, Heiko A Schmidt, Tamara Drucks, Arndt von Haeseler

## Abstract

Selecting the best model of sequence evolution for a multiple-sequence-alignment (MSA) constitutes the first step of phylogenetic tree reconstruction. Common approaches for inferring nucleotide models typically apply maximum likelihood (ML) methods, with discrimination between models determined by one of several information criteria. This requires tree reconstruction and optimisation which can be computationally expensive. We demonstrate that neural networks can be used to perform model selection, without the need to reconstruct trees, optimise parameters, or calculate likelihoods.

We introduce ModelRevelator, a model selection tool underpinned by two deep neural networks. The first neural network, NNmodelfind, recommends one of six commonly used models of sequence evolution, ranging in complexity from Jukes and Cantor to General Time Reversible. The second, NNalphafind, recommends whether or not a *Γ*--distributed rate heterogeneous model should be incorporated, and if so, provides an estimate of the shape parameter, α. Users can simply input an MSA into ModelRevelator, and swiftly receive output recommending the evolutionary model, inclusive of the presence or absence of rate heterogeneity, and an estimate of α.

We show that ModelRevelator performs comparably with likelihood-based methods and the recently published machine learning method ModelTeller over a wide range of parameter settings, with significant potential savings in computational effort. Further, we show that this performance is not restricted to the alignments on which the networks were trained, but is maintained even on unseen empirical data. We expect that ModelRevelator will provide a valuable alternative for phylogeneticists, especially where traditional methods of model selection are computationally prohibitive.

## Introduction

Modelling the process of sequence evolution is a necessary step in carrying out phylogenetic inference. Jukes and Cantor (Jukes and Cantor 1969) introduced the first model of sequence evolution (JC), based on the assumptions that frequency of bases, and pairwise substitutions between each of them, were equally likely. Since then modelling the evolutionary process has been an area of continual and ongoing development and hundreds of models are now available to choose from (K2P (Kimura 1980), F81 (Felsenstein 1981), HKY (Hasegawa et al. 1985), TN93 (Tamura and Nei 1993), and GTR (Tavaré and Others 1986) to name a few).

Systematically discriminating between the available models is a problem that has been approached in a variety of ways, including likelihood ratio tests, Bayes factors, and cross validation. Endorsed by Posada and Buckley (Posada and Buckley 2004), the most common current approach is to carry out maximum likelihood (ML) inference under a selection of different models, then use an information criterion such as Akaike’s Information Criterion (AIC), or the Bayesian Information Criterion (BIC) (Posada and Crandall 1998; Johnson and Omland 2004; Posada and Buckley 2004; Abascal et al. 2005; Posada 2008; Darriba et al. 2012; Kalyaanamoorthy et al. 2017). Model selection software such as ModelFinder (Kalyaanamoorthy et al. 2017) automate the process to a large extent by evaluating models based on BIC (by default), and are employed as a matter of course as part of most phylogenetic analyses. However, there is considerable discussion within the literature about the suitability of information criteria as a discriminating tool within phylogenetics (Grievink et al. 2010; Jhwueng et al. 2014; Seo and Thorne 2018; Susko and Roger 2020; Crotty and Holland 2022). Many of the concerns raised in these articles amount to foundational assumptions of information-theory-based model selection that are necessarily violated within a phylogenetic inference framework. Unburdened by these foundational assumptions, a machine learning approach to the problem is therefore immune to the inherent theoretical complications of the information theory approach.

Theoretical shortcomings aside, the rapid expansion of the size of multiple sequence alignments (MSAs) typically available to empiricists means the computational cost of traditional model selection methods is becoming increasingly prohibitive. Large phylogenomic alignments, consisting of many concatenated genes, should not be assumed to have evolved homogeneously. To address this, different models are often assigned to different sections of the alignment (typically genes or codon positions). Methods like PartitionFinder (Lanfear et al. 2012; Frandsen et al. 2015) accomplish this, but they require the repeated estimation of the appropriate model, sometimes across thousands of genes (Faircloth et al. 2012; Lemmon et al. 2012; Jombart et al. 2014). Thus, model selection can become a computational bottleneck.

Although machine learning methods have found broad application in biology (Tarca et al. 2007; Kandoi et al. 2015; Leung et al. 2016; Kan 2017; Silva et al. 2019), within the field of phylogenetics, they have thus far only been applied to a very limited extent (Tao et al. 2019; Leuchtenberger et al. 2020; Suvorov et al. 2020; Zou et al. 2020). Further refining the scope to model selection within phylogenetics, ModelTeller (Abadi et al. 2020), which utilises a random forest-based machine learning approach, is the only current contribution, although it does not focus primarily on model selection.

To address these issues, we have developed ModelRevelator, a machine learning approach to the model selection problem that is based on neural networks with many layers (deep learning) (LeCun et al. 2015), with the focus on finding the best model of sequence evolution. Further, in the case that the alignment is best modelled incorporating a *Γ*-distributed rates across sites component (Yang 1994), ModelRevelator provides an estimate of the shape parameter. To show the applicability of deep learning in phylogenomics, we compare the results offered by ModelRevelator to ModelFinder (which is representative of current standard practice) as implemented in IQ-Tree (Nguyen et al. 2015; Minh et al. 2020). Using simulated alignments, we assessed the performance of the two methods based on how often they correctly identified the generative model, as well as the accuracy of topological inference carried out using their recommendations. We find that ModelRevelator can estimate the model of sequence evolution as well as the parameter *α* at a comparable accuracy to ModelFinder, with potential for significant reduction in computational expense.

## Methods

### Empirical Lanfear dataset

For testing and evaluation of the neural networks and ML+BIC on empirical data, we used a database of MSAs, collected from the literature by Rob Lanfear, and available at https://github.com/roblanf/BenchmarkAlignments. This collection consists of protein and nucleotide MSAs, from which we selected alignments consisting of DNA alignments of multiple genes. These alignments originated from 31 publications, a full list of which can be found in the supplementary material. We split the 31 multi-gene alignments by gene, resulting in 1,843 individual loci alignments, ranging from 3 to 4,836 taxa. Table 1 shows the distribution of trees with different taxa, trees with 3 to 100 and 101 to 200 taxa being the most frequent trees in the dataset. The length of the individual alignments range from 12bp to 11,049bp with a median length of 261bp (see Table 2).

**Table 1:** Distribution of tree sizes in the empirical Lanfear dataset. Histogram with 200 bins. Column ‘# of taxa’ indicates the range of a tree size bin, column ‘# of trees in bin’ indicates the number of trees in each bin.

| # of taxa | # of trees in bin |
| --- | --- |
| 3 - 100 | 877 |
| 101 - 200 | 910 |
| 201 - 300 | 31 |
| 301 - 400 | 2 |
| 401 - 500 | 4 |
| 501 - 600 | 8 |
| 601 - 700 | 1 |
| 701 - 1000 | 1 |
| 1001 - 2000 | 4 |
| 2001 - 5000 | 5 |

**Table 2:** Distribution of MSA length of original Lanfear MSAs. Bins for histogram = 200, Median length: 261bp, Minimum length = 12bp, Maximum length = 11,049bp. Column ‘# of bp indicates the range of MSA length bin, column ‘# of MSAs in bin’ indicates the number of MSAs in each bin.

| # of bp | # of MSAs |
| --- | --- |
| 12-100 | 50 |
| 101-200 | 483 |
| 201-300 | 548 |
| 301-400 | 231 |
| 401-500 | 166 |
| 501-600 | 91 |
| 601-700 | 75 |
| 701-1000 | 100 |
| 1001-2000 | 64 |
| 2001-4000 | 16 |
| 4001-12000 | 3 |

We used stable IQ-Tree release 1.6.12 to carry out ML inference on the 1,843 loci alignments under a GTR model of evolution. From the results of these analyses we constructed distributions for the five relative substitution rate parameters (G<->T is fixed to a value of 1, with the remaining rates expressed relatively), the empirical base frequency parameters, and the internal and external edge length parameters. We fit splines to these distributions using the AstroML Python package (VanderPlas et al. 2012), as shown in Supplementary Figures 1-3.

### Simulation of training and test datasets

We simulated all MSAs used for training and testing the neural networks using Seq-Gen (Rambaut and Grassly 1997). We generated alignments using six models of sequence evolution with continuous *Γ*-distributed rate heterogeneity across sites (Rhet): JC+G, K2P+G, F81+G, HKY+G, TN93+G, and GTR+G; and their rate-homogeneous (Rhom) versions: JC, K2P, F81, HKY, TN93 and GTR. We ensured that an equal number of training alignments were simulated for each of the 12 models of sequence evolution considered.

We simulated trees by generating a random tree topology with 8, 16, 64, or 128 taxa. We then added internal and external edge lengths drawn randomly from the edge length distributions constructed from the Lanfear data. Depending on which model of sequence evolution was being used to generate the MSA, we drew substitution rate and base frequency parameters as required, from the appropriate Lanfear-based parameter distributions. For NNmodelfind, all training MSAs were of length 1kbp. We also tried using 10kbp MSAs for training NNmodelfind but this did not result in better performance compared to 1kbp MSA training data. For NNalphafind, all training MSAs were of length 10kbp.

For simulating the 6 Rhet models (JC+G, K2P+G, K81+G, HKY+G, TN93+G and GTR+G), we used a continuous *Γ*-distributed rate heterogeneity model of Seq-Gen with 17 distinct alpha parameter values (0.001, 0.01, 0.05, 0.1, 0.3, 0.5, 0.7, 1, 2, 3, 4, 5, 6, 7, 8, 9 and 10) (using the -a command line parameter). We generated an equal number of MSAs for each of those individual alpha parameters.

To test NNmodelfind, NNalphafind, as well as ML approaches, a common test dataset was created. This test dataset consisted of 8, 16, 64 and 128 taxa MSAs, and sequence lengths of 100bp, 1kbp, 10kbp and 100kbp. For the 1kbp test dataset, we also generated MSAs with 256 and 1,024 taxa. For all combinations of taxa and sequence lengths, 512 MSAs were generated for each Rhom model. For Rhet models, we generated 2,048 MSAs per Rhet model and sequence length combination. These 2,048 MSAs were then divided among the 4 taxa levels and then randomly distributed among the 17 distinct alpha parameter values, yielding, on average, 30 MSAs per taxa level and distinct alpha parameter combination. This resulted in a total of 98,304 MSAs in our test dataset.

### Model selection, alpha estimation and tree reconstruction by ML

Two different analyses have been performed using the classical ML approach, namely model selection and parameter estimation.

For model selection, we ran ModelFinder as implemented in stable IQ-Tree release 1.6.12 but restricted the pool of available models to the 12 models on which the networks were trained: JC, K2P, F81, HKY, TN93, GTR, JC+G, K2P+G, F81+G, HKY+G, TN93+G, and GTR+G (using options -mset JC,K2P,F81,HKY,TN,GTR and -mrate E,G). Furthermore, we avoided any identical sequences to be discarded from the alignment using option -keep-ident. To infer rate heterogeneity, we used discrete Γ-distribution approximation with 4 rate categories which is the default in IQ-Tree for Γ models.

As the decision criterion, we used BIC as is default in IQ-Tree. To explicitly estimate the alpha parameter, we ran ModelFinder for each alignment, but restricted the pool to include only the heterogeneous models. We then took the reported alpha value from the best heterogeneous model as our estimate of alpha.

According to the IQ-Tree documentation, IQ-Tree applies a lower limit of 0.02 for the shape parameter alpha. To allow the estimation of lower alpha values we reset this limit to 0.0001 using the command line flag -amin 0.0001. For some alignments there were errors caused by numerical issues especially for very low alpha values. We resolved these by re-running with the -safe option as suggested by the error message of IQ-Tree.

In order to reconstruct trees for comparison, we ran IQ-Tree independently with the model of sequence evolution chosen by ModelRevelator and ML+BIC respectively. If the model chosen by ModelRevelator included heterogeneous rates then the alpha parameter was fixed to the recommended value. If the model chosen by ML+BIC included heterogeneous rates then the alpha parameter was optimised in the tree reconstruction process.

### Data preprocessing for NNs

To be able to use the variably-sized simulated MSAs as input for training and testing of NNmodelfind and NNalphafind, we converted each alignment into a format of fixed-sized.

For NNmodelfind, we used summary statistics of sequence pairs of an MSA. To maximise the information in the input, we decided to use 10,000 randomly drawn sequence pairs, ensuring that the same sequence was not chosen for both sequences of a summary statistic to be calculated. These 10,000 summary statistics consisted of 26 features each (substitution counts: A-C, A-G, A-T, C-G, C-T, and G-T, in both directions = 12 features; invariant site counts: A-A, C-C, G-G, T-T = 4 features; 4 nucleotide counts per sequence = 8 features; total transition count and total transversion count = 2 features). This yields a total feature count for each MSA of 10,000 × 26. All of these summary statistics are normalised by the sequence length of the MSA. For more efficient processing of the convolution layers of the ResNet-18, we reshaped these input features into an input tensor of shape 40 × 250 × 26.

For NNalphafind, we generated normalised base composition profiles of 10,000 sites and the 4 possible bases, yielding an input size of 10,000 × 4. We also experimented with summary statistics incorporating column-wise transitions and transversion, however this yielded worse performance than just base counts. For both networks, for all MSAs which were smaller or larger than 10,000 positions, we randomly over- or undersampled the MSA to achieve the required dimension for the input.

### NNmodelfind architecture

Figure 1 shows the architecture of NNmodelfind. It is based on a ResNet-18 architecture (He et al. 2015a), but with an adapted input strategy. The input, as described above, has dimensions 40 × 250 × 26. We replaced the initial pooling layer of a ResNet-18 with 4 encoding 2D convolution layers, with 2×1 kernels and 32, 64, 96 and 96 channels. The output of the last 2D encoding layer is then passed to 4 standard ResNet-18 blocks with 3×3 convolutions and 96, 192, 384 and 768 channels. The output layer has 6 categories, one for each generative model (JC, K2P, K81, HKY, TN93, GTR). The loss function used was a categorical cross entropy loss function. We applied Batch Normalisation (Ioffe and Szegedy 2015) before each ReLu activation function and L2 Regularisation (Cortes et al. 2012), for all convolution layers to improve ResNet-18 training. Additional parameters used were a learning rate of 1×10^-5, Adam optimiser (Kingma and Ba 2014) for gradient descent, He Normal initialisation (He et al. 2015b) for the weights and zeros initialisation for biases.

**Figure 1:**
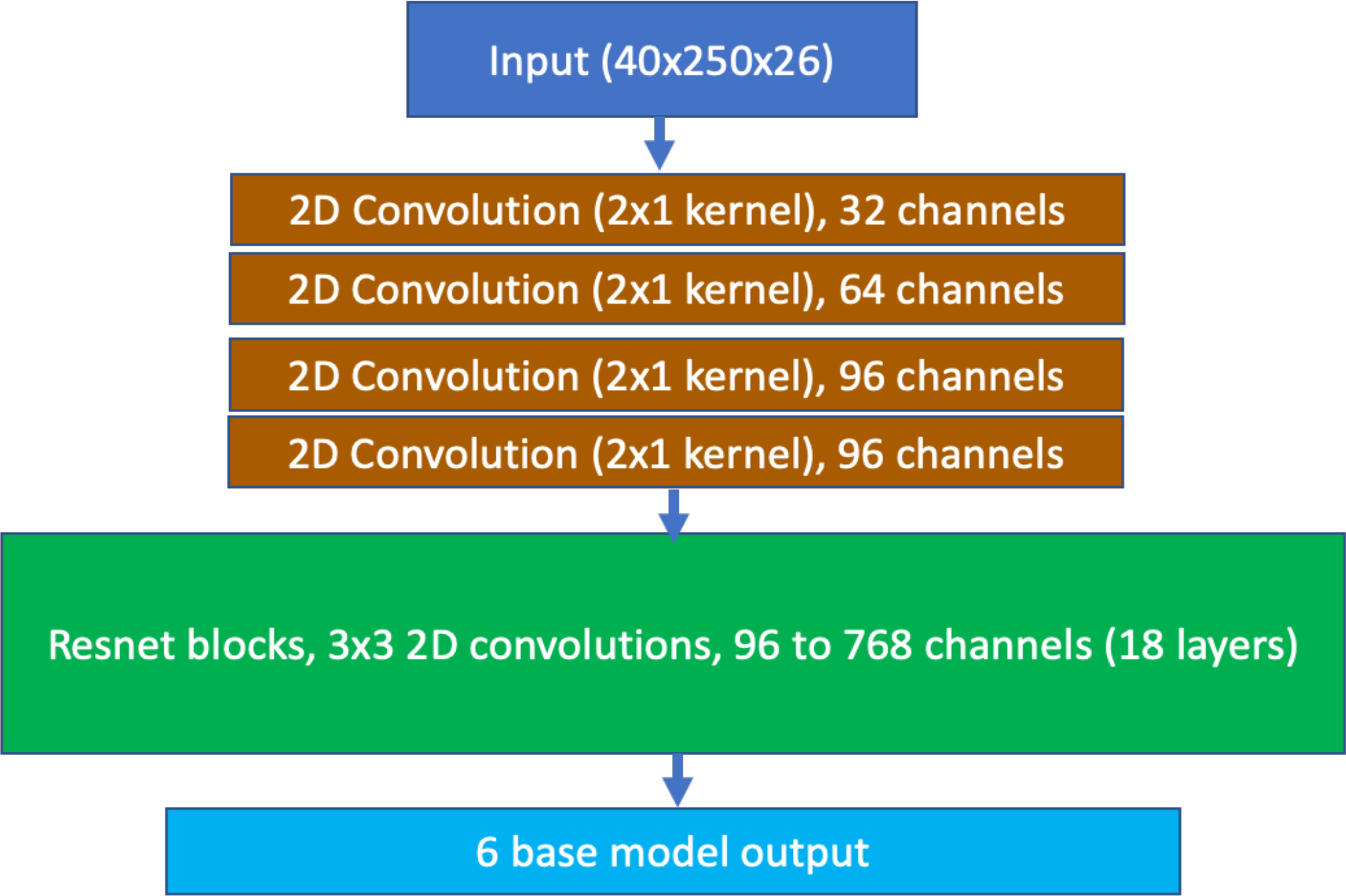
Schematic of NNmodelfind. The architecture follows a ResNet-18, with a modified input strategy.

### NNalphafind architecture

As shown in figure 2, NNalphafind is a combination of 1D convolutions for feature encoding and a bidirectional LSTM with an Attention layer (Raffel and Ellis 2015). The input consists of profiles of 10,000 positions by 4 base frequencies. The 3 layers of 1D convolutions have 256, 512 and 768 channels, followed by a bidirectional LSTM with 1,200 steps, a 1D pooling layer with a pooling window of 4, and an Attention layer with a step dimension of 2,498. NNalphafind has 2 outputs, a categorical output and a scalar output. The categorical output estimates whether an MSA should be modelled with or without rate heterogeneity. The loss function is also a categorical cross entropy loss function. The scalar output estimates the alpha parameter, achieved by a mean absolute percentage error function. Weights of NNalphafind were initialised using Glorot Uniform initialisation (Glorot and Bengio 2010) and biases were initialised with zeros.

**Figure 2:**
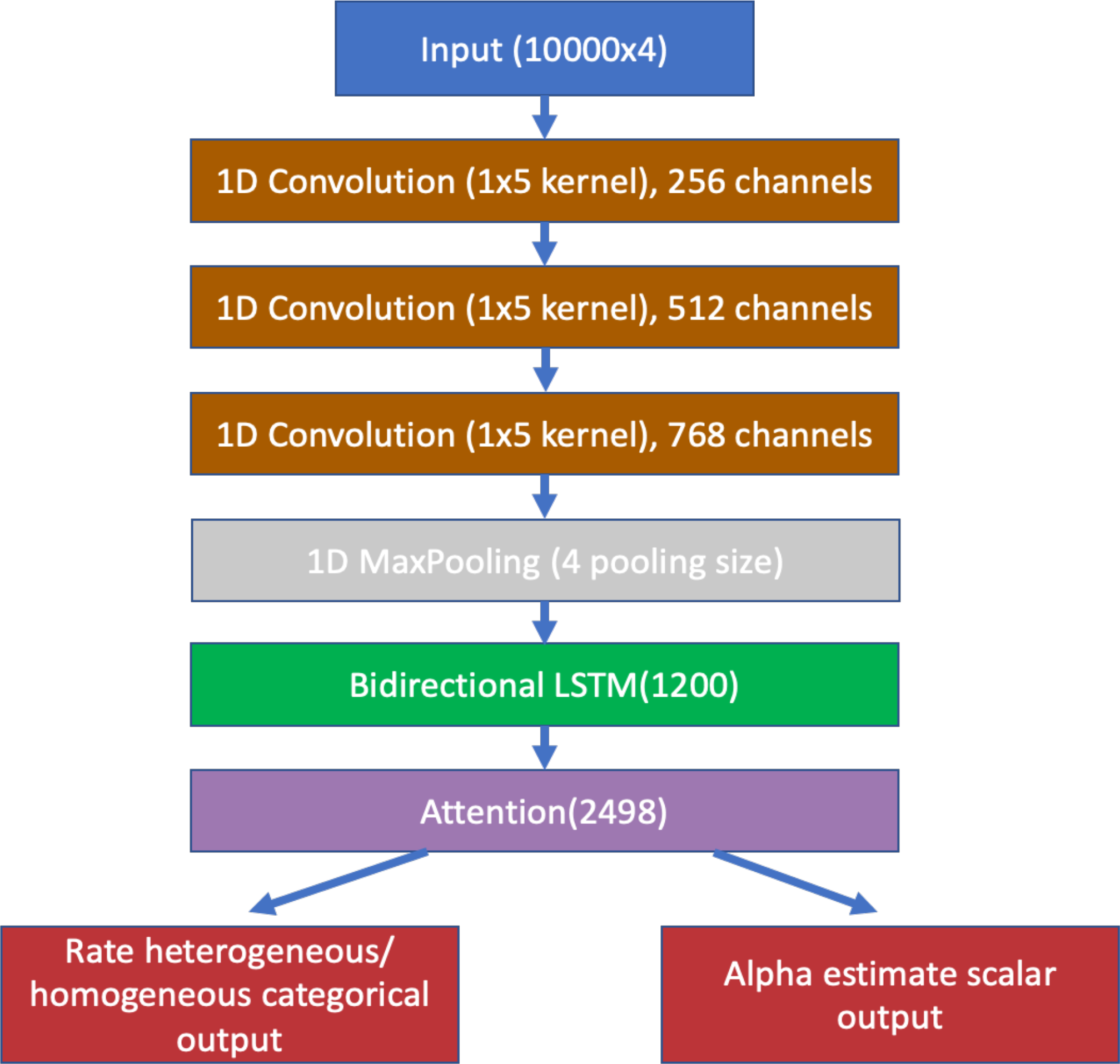
Schematic of NNalphafind.

### Training NNmodelfind and NNalphafind

We trained NNmodelfind, as well as NNalphafind, with the training dataset described above, using a batch size of 40 and a learning rate of 1×10^-5 for both neural networks. For NNmodelfind, we required 76 epochs for convergence and for NNalphafind, we required 341 epochs for convergence. We used Tensorflow 2.4 (Abadi et al. 2016) as a framework for training and testing of our neural networks. As GPUs for training, we used Nvidia Tesla V100 with 32Gb of memory, or Nvidia RTX2080Ti with 11Gb of memory.

To avoid overfitting, we performed early stopping when:

1. there was no improvement in accuracy on the validation dataset for 5 epochs for NNmodelfind and for 15 epochs for NNalphafind; and
2. the divergence of training and validation accuracy was not higher than 4% for NNmodelfind and 6% for NNalphafind.

In total 242,688 MSAs were used to train the networks.

### Testing NNmodelfind and NNalphafind

For testing the neural networks, we used the datasets described earlier and the trained models for NNmodelfind and NNalphafind. Testing was either performed using the Keras predict() function or ONNXRuntime 1.6.0 (https://github.com/microsoft/onnxruntime). ONNXRuntime is a dedicated neural network execution library for the open ONNX neural network exchange format (https://onnx.ai/). In order to use ONNXRuntime, we first exported the Keras/Tensorflow neural network graphs to the ONNX format using the tools keras2onnx and tf2onnx of the Python package onnx 1.9.0.

### Comparing ModelTeller and ModelRevelator

For running ModelTeller, we downloaded the latest version from GitHub (https://github.com/shiranab/ModelTeller, commit hash 897bd6925606416071dcf3c1a03422f8e7a4e7b7; ModelTeller trained models downloaded 2021-06-06) and followed the installation guidelines. Then, we ran ModelTeller on a subset of (100bp, 1kbp and 10kbp MSAs) our common Lanfear test set MSAs, using the -m command line parameter of ModelTeller to just provide the MSA files which should be processed. We then took the top ranked model ModelTeller output for all further analysis. We also used the trees which ModelTeller estimated with PhyML, which is distributed as an integral part of ModelTeller, for comparison with ModelRevelator. The branch score distances as well as the Robinson-Foulds distances were calculated using Treedist of the PHYLIP package (https://evolution.genetics.washington.edu/phylip/).

### Time measurements for ModelRevelator and ML

For the neural networks, we performed all measurements using ONNXRuntime 1.6.0 for CPUs, thus no GPU was used for inference. For comparison to single-threaded IQ-Tree, we forced ONNXRuntime to only run on a single core by using the Linux tool *taskset*. All benchmarks were run on the same machine equipped with two 32 core AMD Epyc 7551 CPUs on the operating system OpenSuSE Linux 15.0.

All IQ-Tree analyses were run on the same hardware as the neural network runtime measurements, using only one CPU core each. All runtime measurements for ModelRevelator as well as ML+BIC were performed using a 10% representative subset of alignments sampled from the test dataset. The reported runtimes are the averages of the measured runtimes.

## Data availability

Python source code for executing ModelRevelator, generating training, validation and test datasets and the Tensorflow implementations of NNmodelfind and NNalphafind of ModelRevelator are available from https://github.com/cibiv/ModelRevelator. NNmodelfind and NNalphafind of ModelRevelator are also available in ONNX format in the same repository. Due to the amount of data (240,000 MSAs of training data, 170,000 MSAs of test data, 603 GB in total), the MSAs in phylip format and the corresponding tree files from the common test dataset are available upon request from the authors.

ModelRevelator will also be made freely available as part of the forthcoming version of IQ-Tree (http://www.iqtree.org),

## Results

We introduce ModelRevelator, a machine learning-based approach to model selection for phylogenetic inference. Users can input their MSAs to ModelRevelator. ModelRevelator will then advise which model of sequence evolution should be used, whether a *Γ*-distributed rate heterogeneity component should be included, and if so, an estimate of the shape parameter. ModelRevelator consists of two neural networks which operate independently to provide model recommendations that can be adopted to significantly accelerate inference time. The first network, NNmodelfind, was trained to select a model of sequence evolution prior to phylogenetic inference, thereby bypassing the computationally expensive procedure of performing model selection via ML and information criteria. The second network, NNalphafind, was trained to make recommendations with respect to rate heterogeneity, and its output is two-fold. NNalphafind will first recommend whether the alignment is characteristic of substitution rate heterogeneity, or whether a rate homogeneous model will suffice. If the network recommends that a *Γ*-distributed rates across sites model is appropriate, it will then provide an estimate of the shape parameter which can be incorporated into the inference. Both networks were trained on simulated data based on empirical alignments. We show here that the networks have comparable accuracy to traditional model selection methods, and have significant advantage in terms of computational expense.

### Generating testing and training data

In order to train our neural networks a large number of simulated multiple sequence alignments (MSAs) were required. However, we were mindful that ultimately, we require the neural networks to be useful tools for the analysis of empirical data, and thus we needed our training MSAs to be broadly representative of empirical datasets. To this end we obtained a large database of empirical MSAs made available to us by the Lanfear group of Australian National University (https://github.com/roblanf/BenchmarkAlignments). From the Lanfear MSAs we reconstructed trees via maximum likelihood inference with the stable IQ-Tree release 1.6.12, under six different models of sequence evolution, with and without accounting for rate heterogeneity. We thus obtained empirical distributions for internal edge lengths, external edge lengths, substitution rates, and base frequency parameters. We fit cubic splines to these empirical distributions, enabling us to sample random values as required.

We then generated random tree topologies of varying sizes, and allocated edge lengths to these topologies by randomly sampling from the external and internal edge distributions. We also constructed models of sequence evolution by randomly sampling from the substitution rate and base frequency distributions. These tree topologies and substitution models were then used as input to Seq-Gen (Rambaut and Grassly 1997), in order to generate MSAs of 1kbp length for training of the neural networks. We also generated MSAs incorporating the continuous-Γ model of rate heterogeneity, with alpha parameters ranging from 0.001 (extreme rate heterogeneity) up to 10 (mild rate heterogeneity). Testing data was generated in essentially the same way, although we tested a wider range of alignment sizes, ranging from 100bp up to 100kbp.

Figure 3 illustrates the procedure we adopted for generating alignments. A fuller account of the precise parameter settings and volume of MSAs generated can be found in the methods section.

**Figure 3:**
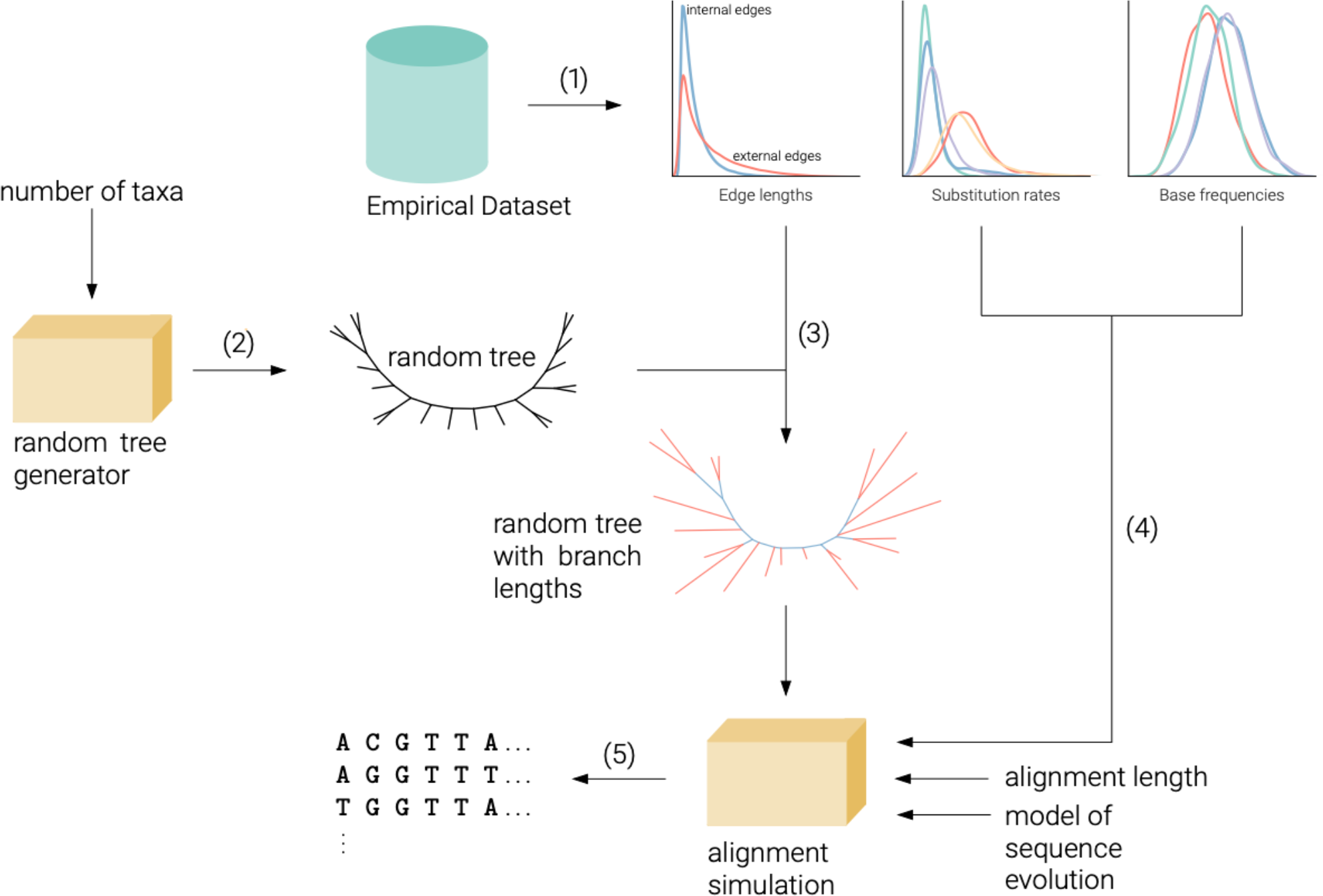
Schematic illustration of the workflow followed to produce training and testing data for the neural networks. (1) Edge lengths, substitution rates, and base frequencies are inferred using maximum likelihood for 1,842 empirical MSAs, to form empirical distributions of these parameters. (2) For the specified number of taxa a random tree topology is generated following a Yule process. (3) Edge lengths for the tree are drawn randomly from the distribution of internal and external edges obtained in (1). (4) For a given substitution model, the appropriate model parameters are drawn from the distributions obtained in (1). (5) For a given alignment length an MSA is simulated based on the tree and model of sequence evolution obtained from (3) and (4) respectively, using the program seq-gen.

### Estimating the evolutionary model

Our first objective was to train a neural network, which we have called NNmodelfind, to successfully estimate the correct model of sequence evolution. As described in the methods section, we trained NNmodelfind on a wide variety of simulated MSAs. The training MSAs were generated according to one of 6 models of sequence evolution (JC, K2P, F81, HKY, TN93, and GTR). We incorporated an equal number of MSAs that were simulated under rate homogeneous (Rhom) and rate heterogeneous (Rhet) conditions. Rate heterogeneous alignments were simulated with the shape parameter, alpha, taking one of 17 potential values between 0.001 and 10. MSA size was either 8, 16, 64, or 128 taxa, with each size equally represented in the training data. The length of all training MSAs was 1kbp. Experiments using training data of 10kbp length did not show improvements in NNmodelfind accuracy over 1kbp training data.

Having trained NNmodelfind, we generated test MSAs to evaluate its performance. Test MSAs were generated using the same parameter settings as the training MSAs, with additional MSAs generated with higher numbers of taxa (256, and 1024), and different sequence lengths (100bp, 10kbp, and 100kbp) in order to evaluate the generalisability of NNmodelfind.

By way of comparison we used a traditional method of model selection via maximum likelihood inference to choose the model of sequence evolution for each test MSA. We used the ModelFinder approach within IQ-Tree, with BIC set as the discriminating criterion. We refer to this method of model selection as ML+BIC.

Figure 4 shows the accuracy for Rhom test MSAs of length 1kbp. Figure 4a displays the accuracies separately for the individual models and taxa levels. Figure 4b shows confusion matrices for each method and taxa level, detailing the distribution of the inferred model for each true model. Figure 5 shows the analogous information to Figure 4, for Rhet test MSAs. Overall, Figures 4 and 5 indicate that NNmodelfind and ML+BIC are able to estimate the generative model of sequence evolution with similar accuracy. There are modest differences between the two methods, for example ML+BIC is more accurate when the true model is GTR or TN93, whereas NNmodelfind is more accurate when the true model is F81 or HKY.

**Figure 4:**
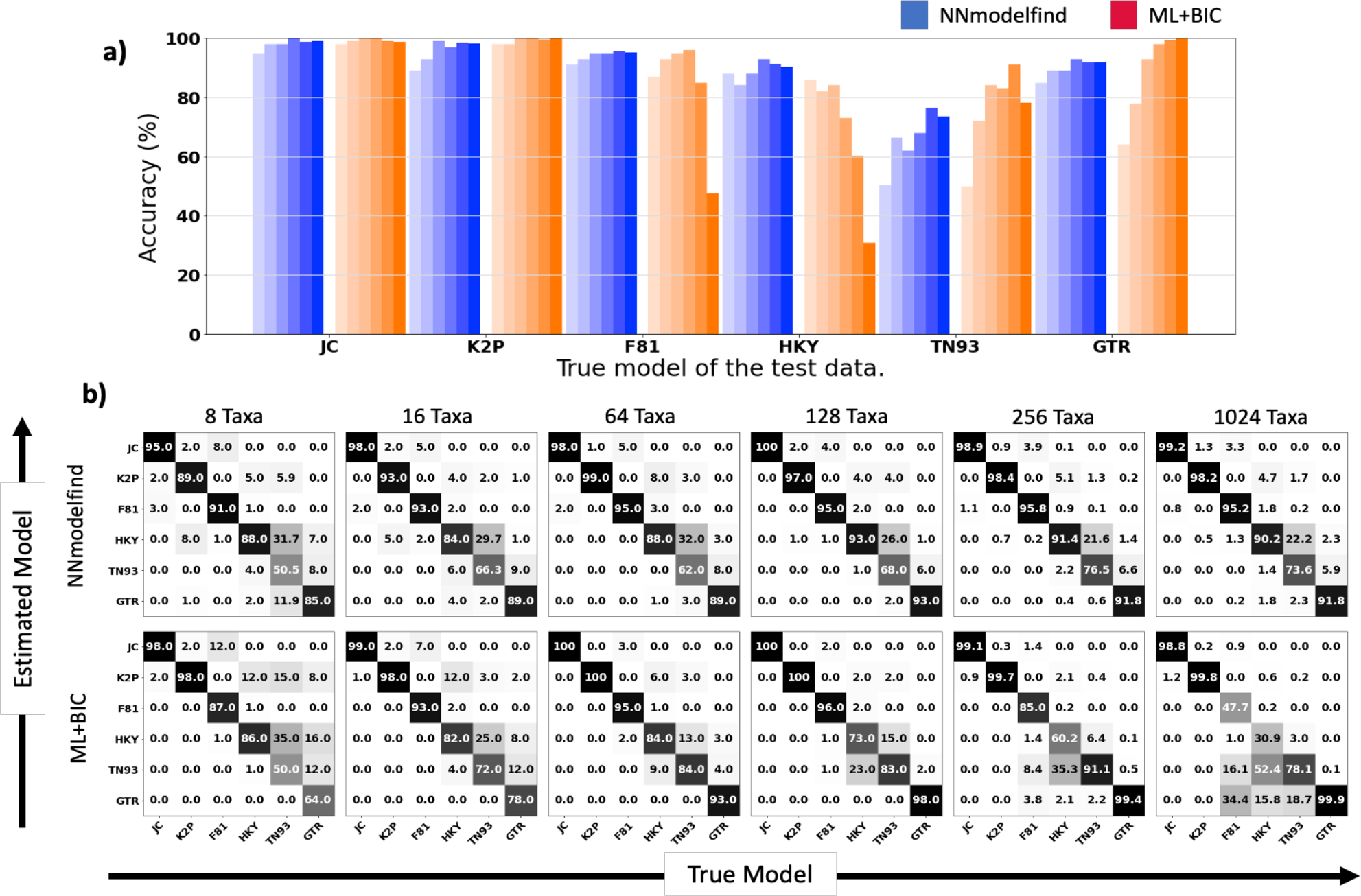
Model selection accuracy of NNmodelfind and ML+BIC on 1,000bp long Rhom MSAs. **(A)** Blueish colours indicate results from NNmodelfind, reddish colours indicate results from ML+BIC. Darkening bars distinguish increasing number of taxa from left to right (8, 16, 64, 128, 256 and 1024). **(B)** Confusion matrices for NNmodelfind and ML+BIC. Entries along the diagonal indicate the percentage of alignments for which the correct model was identified.

**Figure 5:**
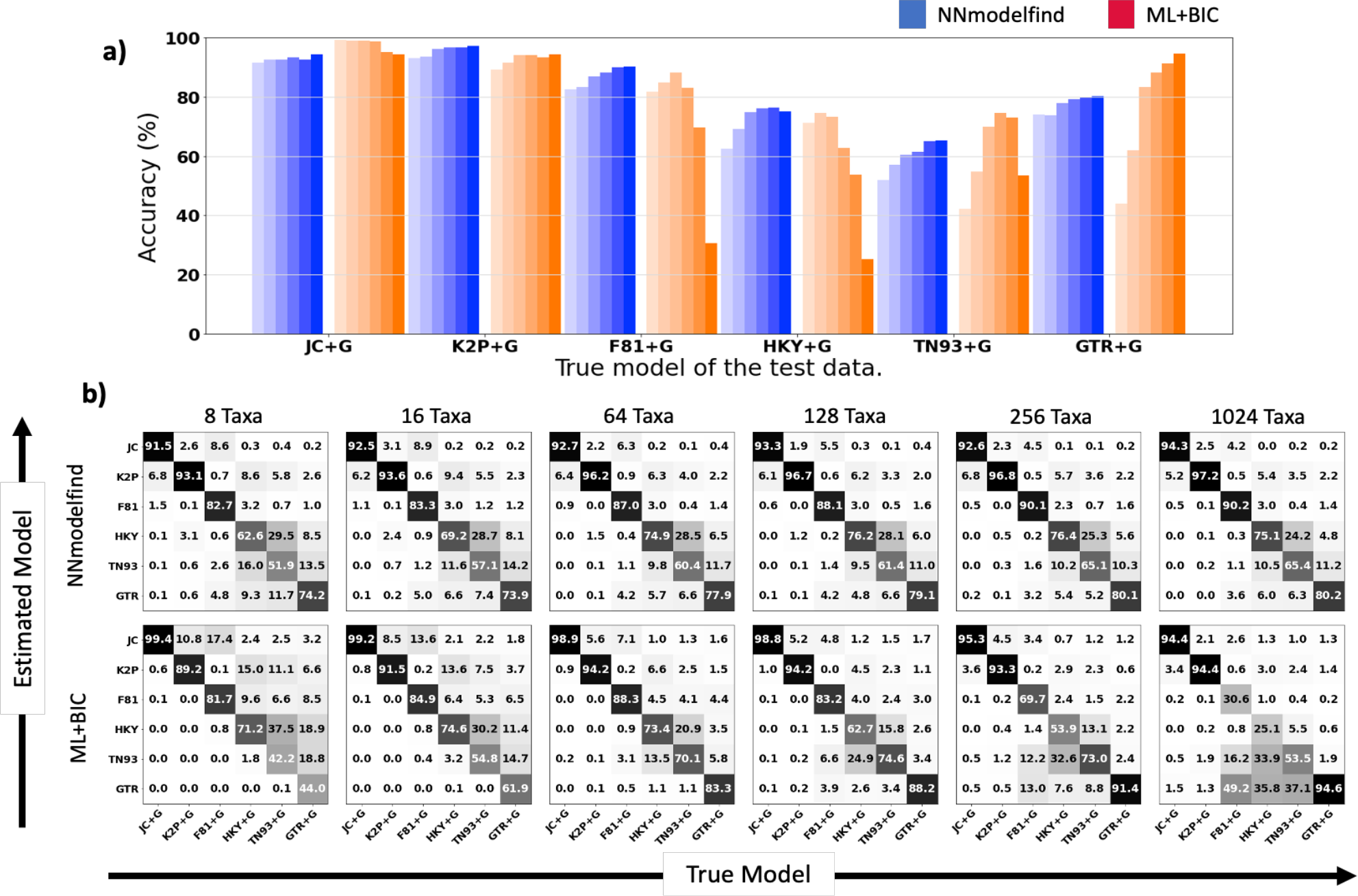
Model selection accuracy of NNmodelfind and ML+BIC on 1,000bp long, Rhet MSAs. **(A)** Blueish colours indicate results from NNmodelfind, reddish colours indicate results from ML+BIC. Darkening bars distinguish increasing number of taxa from left to right (8, 16, 64, 128, 256 and 1024). **(B)** Confusion matrices for NNmodelfind and ML+BIC. Entries along the diagonal indicate the percentage of alignments for which the correct model was identified.

Figures 4b and 5b show the confusion matrices associated with the 1kbp test alignments. The confusion matrices reveal an interesting phenomenon in relation to the performance of ML+BIC. For low numbers of taxa, when ML+BIC infers the incorrect model, it almost exclusively selects a model of lower complexity than the true model. Conversely, when the MSAs contain a high number of taxa, ML+BIC tends to err in the direction of more complex models. This is likely a result of the fact that as the number of taxa increases, the log likelihood scores grow and the influence of the penalty term in the BIC formula is reduced. Thus ML+BIC tends to more complex models as the number of taxa increases. By way of comparison, the error structure displayed by NNmodelfind does not show any strong pattern of preference for models of lower or higher complexity, and this appears to be the case independent of the number of taxa in the alignment. This is likely a result of the fact that the input size of NNmodelfind is fixed and so increasing the number of taxa does not influence the error structure.

### Generalisation of NNmodelfind

Although NNmodelfind was trained strictly on MSAs of length 1,000bp, if it is to be of use to empiricists then it must be reasonably generalisable on MSAs of various lengths. To investigate its capabilities in this regard we generated test alignments that were both shorter (100bp) and longer (10kbp and 100kbp) than the training alignments. Supplementary Figures 4 - 9 show the performance of NNmodelfind and ML+BIC for these data, in the same format as Figures 4 and 5.

For short sequences of 100bp, the performance of NNmodelfind is poor for all models except GTR. The confusion matrices suggest that NNmodelfind selects GTR for most MSAs, independent of the model used to generate the alignment. Conversely, ML+BIC performs poorly for more complex models such as GTR, but is very reliable for simpler models such as JC and K2P. The confusion matrices suggest that ML+BIC has a tendency to select a simple model regardless of the model used to generate the alignment. The poor performance of both methods is likely a reflection of the high amount of stochastic noise within the short datasets, and is not unexpected. The performance of both ML+BIC for long alignments of 10kbp and 100kbp was excellent. Supplementary Figures 8 and 9 show that for 100kbp MSAs, ML+BIC identified the generative model with accuracies approaching 100%, across all taxa and model combinations. This is not unexpected, as ML is known to be statistically consistent (Truszkowski and Goldman 2016), and we would therefore expect ML+BIC to perform well for very long sequences.

NNmodelfind however saw no appreciable improvement for these longer sequences. The rate at which NNmodelfind correctly inferred the generative model was similar for 10kbp and 100kbp as it was for 1,000bp. One obvious explanation for this observation is that, irrespective of the actual length of the MSA, the input size for NNmodelfind is fixed. Thus, for long alignments, NNmodelfind samples sites from the alignment up to the required input size, and therefore does not make full use of all the information contained within the alignment.

In addition to generalising over a wide range of sequence lengths, the performance of NNmodelfind on varying numbers of taxa is also of interest. NNmodelfind was trained on MSAs which ranged in size from 8 to 128 taxa. To investigate whether the performance of NNmodelfind could be extrapolated to larger datasets, we simulated test MSAs with 256 and 1024 taxa, using the parameters estimated from the Lanfear MSAs as before. The results are displayed in Figures 4 and 5. Figures 4a and 5a show that the performance of NNmodelfind is not compromised by increasing the number of taxa beyond that used to train the network. For all generative models, NNmodelfind identified the correct model at rates as good as or slightly higher than for the smaller MSAs. For ML+BIC however, increasing taxa had a detrimental effect on the success rate when the generative model was F81, HKY, and to a lesser extent TN93, both with and without rate heterogeneity. The confusion matrices in Figures 4b and 5b suggest that with these generative models, ML+BIC tended to select more complex models, most often choosing GTR.

### Estimating the presence and degree of rate heterogeneity

The assumption that all sites in an MSA mutate at the same rate has long been regarded as biologically implausible. Rate heterogeneity has become an essential component of models of sequence evolution, and is typically modeled by the discrete *Γ* distribution (Yang 1994). The alpha parameter of the *Γ* distribution determines its shape and thus the extent of rate heterogeneity. In conjunction with NNmodelfind, a neural network which can accurately estimate the alpha parameter could further expedite phylogenetic reconstruction. We trained a second neural network, NNalphafind, to first estimate whether a rate homogeneous model was appropriate for the MSA, and if not, to then estimate the alpha parameter of the *Γ*-distributed rate heterogeneity component.

As described in the methods section, NNalphafind was trained on MSAs containing 8, 16, 64 and 128 taxa. Each training MSA was 10kbp long, as opposed to the 1kbp MSAs that were used to train NNmodelfind. NNalphafind was tested on the same test MSAs simulated to test NNmodelfind, but for clarity we reiterate the simulation procedure here with the added detail of the alpha parameter levels. The generative models of sequence evolution used were JC, K2P, F81, HKY, TN93, and GTR. The amount of rate heterogeneity in the MSAs was varied by controlling the rate parameter, alpha. In total there were 17 levels of rate heterogeneity, represented by 17 distinct alpha values (0.001, 0.01, 0.05, 0.1, 0.3, 0.5, 0.7, 1, 2, 3, 4, 5, 6, 7, 8, 9 and 10). Small values of alpha correspond to a high degree of rate heterogeneity, whereas large values of alpha correspond to a low degree of rate heterogeneity. With 4 taxa levels, 6 generative models, 4 sequence length levels, and 17 levels of rate heterogeneity, there were 4 × 6 × 4 × 17 = 1632 parameter combinations. For each of these parameter combinations we simulated 30 test MSAs.

Preliminary analysis suggested that there was no relationship between the generative model of sequence evolution and the accuracy of the alpha inference. Therefore, for simplicity, we did not stratify the results by generative model. This meant we had 1632 / 6 = 272 parameter combinations of interest, with each parameter combination having 180 (30 MSAs for each of the 6 different generative models) test MSAs available to evaluate performance. For each of the 272 parameter combinations we calculated the mean alpha value inferred by NNalphafind, and also by ML+BIC. Given the range of true alpha values spanned several orders of magnitude, to enable a simple comparison we calculated the ratio of the mean inferred alpha values and the true alpha values. A ratio of 1 would therefore correspond to perfect inference (on average), while scores above or below 1 would correspond respectively to an overestimation or underestimation of the true alpha. The performance of the two methods at estimating the true value of alpha in the generative models are summarised in heatmap form in Figure 6. The colour scheme of the heatmap distinguishes between good inference (white), underestimation (grey), and overestimation (red). In general, the lighter the shading the better the inference of alpha.

**Figure 6:**
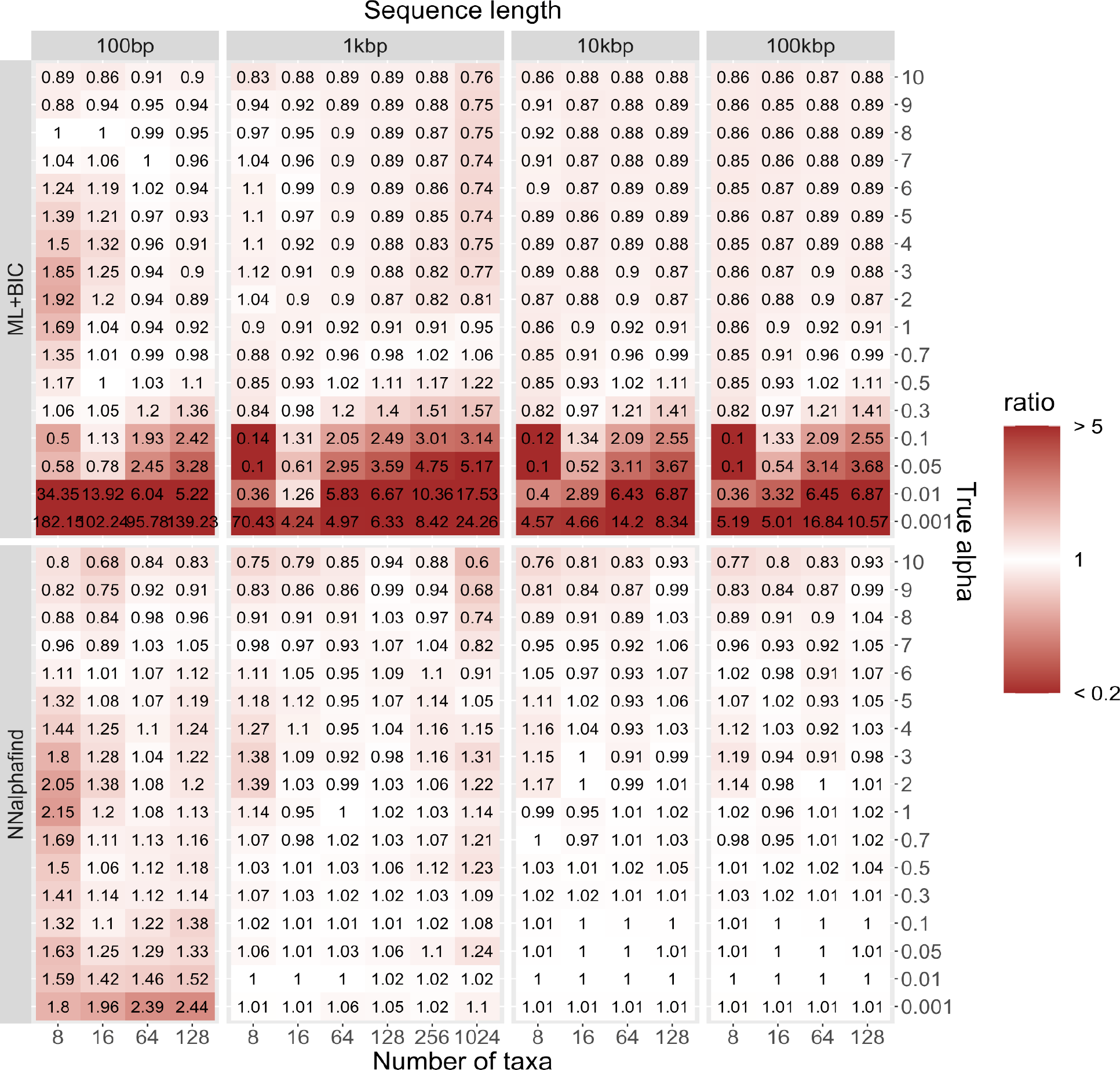
Ratio of mean inferred alpha value and true alpha value, stratified by inference method (ML+BIC or NNalphafind), number of taxa (8, 16, 64, and 128), sequence length (100bp, 1kbp, 10kbp, and 100kbp), and true alpha value (17 levels ranging from 0.001 (very strong heterogeneity) up to 10 (very weak heterogeneity)). Values close to 1 indicate accurate inference. Colour scale indicates degree of bias in estimation. A ratio of 5 indicates a five-fold overestimation of alpha, a ratio of 0.2 indicates a five-fold underestimation of alpha.

Figure 6 indicates that NNalphafind generally performs better than ML+BIC at estimating the true alpha value used to generate the data. NNalphafind was able to accurately infer the true alpha, particularly when alpha was less than 1, indicating strong rate heterogeneity.

Conversely, ML+BIC performs particularly poorly for low alpha values (< 0.3) and performance in this range is not strongly improved with increasing sequence length. Interestingly, the performance of NNalphafind does clearly improve as sequence length increases, in contrast to the performance of NNmodelfind. This is likely due to the fact that the input size of NNalphafind is larger, and the network was trained on 10kbp alignments, as opposed to 1kbp alignments for NNmodelfind. As such NNalphafind has the capacity to utilise the additional information in the longer test MSAs.

### Generalisation of NNalphafind

As with NNmodelfind, the results indicate that NNalphafind also generalises well to sequence lengths that it was not trained on. NNalphafind was trained on 10kbp alignments, and although its performance does deteriorate somewhat for 1kbp alignments, and then further for 100bp alignment, this is to be expected considering the comparative lack of information contained within these shorter alignments. The same effect is evident in the performance of ML+BIC and, regardless of sequence length, NNalphafind is always more accurate for low values of alpha, and similarly accurate to ML+BIC for higher alphas. Figure 6 does not indicate any parameter combinations where NNalphafind is clearly inferior to ML+BIC (See supplementary fig. 10 for mean squared error of inferred alphas).

NNalphafind also generalised well to higher numbers of taxa, although perhaps not as well as NNmodelfind. NNmodelfind saw no drop in performance when tested on 256-taxon and 1024-taxon alignments, whereas the accuracy of NNalphafind’s estimates was increasingly compromised with higher taxa. That said, even at high taxa levels NNalphafind outperformed ML+BIC for small alpha values, and was similarly accurate elsewhere.

### Influence of Model Selection on Phylogenetic Inference

The primary goal of phylogenetic inference is to reconstruct accurate trees. In order to compare the performance of ModelRevelator to ML+BIC, we reconstructed trees using the models recommended by the two methods. When reconstructing trees using the output of ModelRevelator, we fixed the alpha parameter (when rate heterogeneity was recommended) to the value estimated by NNalphafind, rather than allowing it to be optimised. For each MSA, we then compared the trees inferred via ModelRevelator and ML+BIC to the tree used to simulate the alignment. For this, we calculated the normalised Robinson-Foulds distances (Robinson and Foulds 1981) between the three trees. Figure 7 shows that when comparing the Robinson-Foulds distances of the simution tree (*T*_SIM_) vs the reconstructed trees (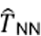 and *T*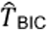 for ModelRevelator and ML+BIC respectively), we observed similar distributions for both methods. Furthermore, Figure 7 also shows that the reconstructed trees are typically much closer to each other than they are to the simulation trees. This effect is most obvious for short sequences, with the difference diminishing as sequence length increases. Thus, reconstructed trees, irrespective of the reconstruction method, appear more in concordance with each other than they are with the generative trees that were used to simulate the alignment, pointing to the same type of error being made by both model estimation methods. An alternative viewpoint would be that both methods are equally hindered by stochastic variation in the simulation process, resulting in inaccurate tree reconstruction, independent of the model chosen by each method. Reinforcing this view, Figure 7 clearly indicates that sequence length is a strong indicator of the accuracy of tree reconstruction.

**Figure 7:**
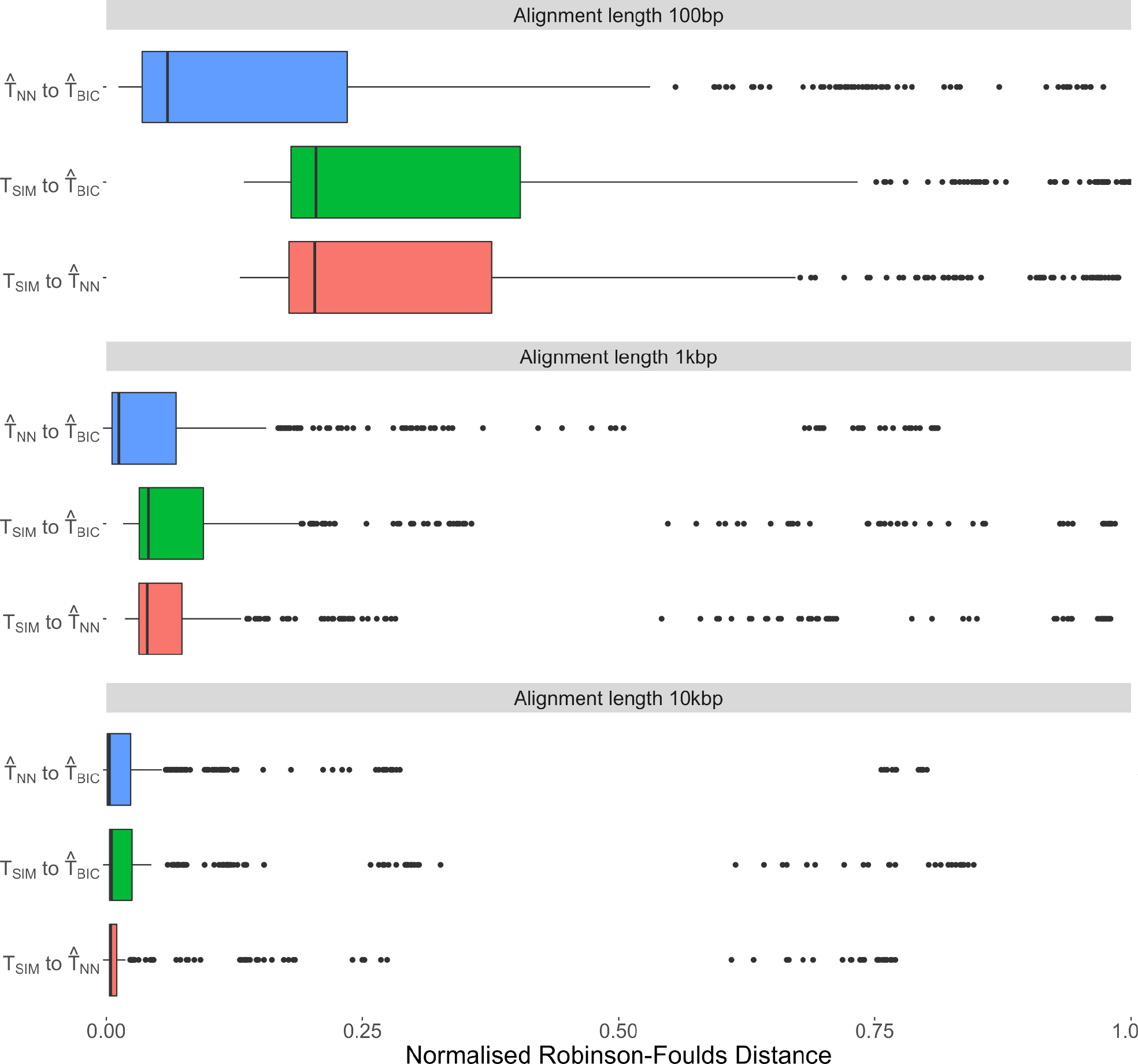
Estimating concordance in tree reconstruction by calculating the normalised Robinson-Foulds distances of trees reconstructed using the model recommended by ModelRevelator 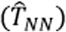 or ML+BIC 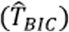, and comparing these results to the trees used to simulate the MSAs (T_SIM_). Results for MSAs of length 100bp, 1kbp and 10kbp are shown in the top, middle and bottom panel, respectively.

### Generalisation to unseen empirical alignments

In addition to generalisation towards large numbers of taxa in an MSA, we also wanted to test generalisation of ModelRevelator to other empirical datasets. This is of obvious practical relevance, but is also important as generalisation of neural networks to potentially differently distributed test datasets is a topic of constant discussion (Ganin and Lempitsky 2015; Sagawa et al. 2019). We used 6,453 MSAs from the original PANDIT dataset (Whelan et al. 2006) to perform parameter estimation and tree reconstruction by ML under a GTR+G model, with IQ-Tree. Supplementary Figures 1 to 3 show the distributions of the edge length, substitution rate, and base frequency parameters for both the Lanfear and PANDIT datasets. The parameters are clearly differently distributed between the two datasets, most notably in regard to the edge lengths.

For each PANDIT MSA, using the inferred tree and model parameters (including the shape parameter, alpha), we then used Seq-Gen to simulate 10 MSAs (with GTR+G as the generative model) of the same length as the PANDIT MSA, creating 64,530 simulated PANDIT-type MSAs in total. On these PANDIT type MSAs we performed model estimation using both ML+BIC and ModelRevelator. Figure 8 shows a concordance matrix for the results of model selection with the two methods. Overall, ML+BIC inferred the correct model of GTR+G in 69% of alignments, compared to just 61% for ModelRevelator. To investigate these results in more depth, we compare the two methods on three measures: (1) the binary determination of whether a rate heterogeneous model should be employed; (2) the decision of which of the six models of sequence evolution should be adopted; and (3) the resulting topological accuracy of trees inferred.

**Figure 8:**
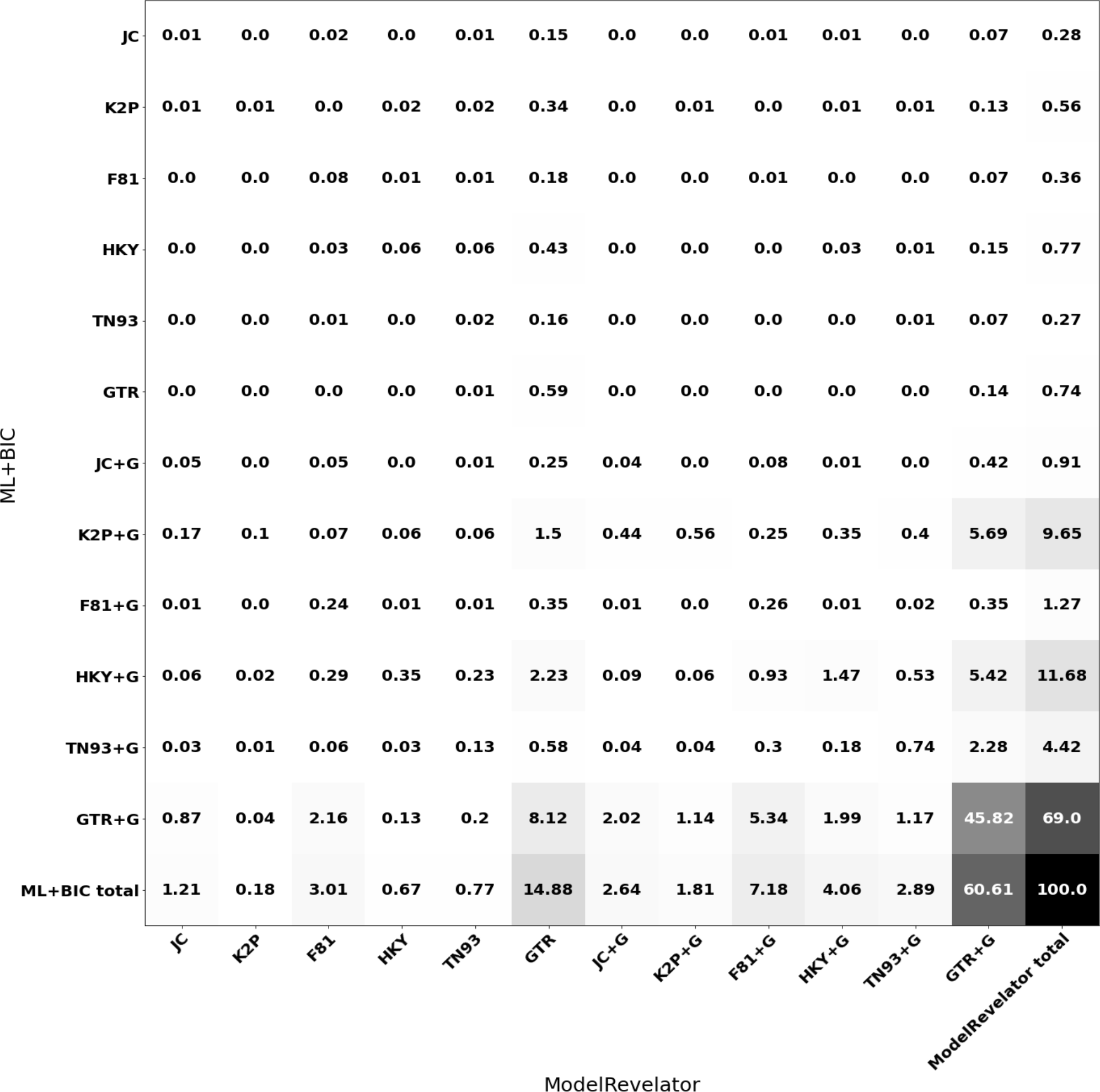
Concordance matrix of ModelRevelator and ML+BIC estimates for MSAs simulated under PANDIT trees and parameters. The generative model for all MSAs was GTR+G. Column and row sums indicate proportion of alignments inferred for each model by ModelRevelator and ML+BIC respectively.

With regard to the recommendation to include a rate heterogeneous component in the model, ML+BIC has performed better than ModelRevelator on these alignments. ML+BIC correctly recommended rate heterogeneity in approximately 97% of alignments, compared to approximately 79% for ModelRevelator. This means ModelRevelator is more likely than ML+BIC to make a Type II error, that is to fail to detect the true presence of rate heterogeneity in the data. It is worth considering here that rate heterogeneity is modelled very efficiently by the *Γ* distribution, at a cost of only one parameter. This means the penalty in the BIC calculation is quite small, but it gives increased flexibility to the model to fit more of the variability in the data (whether that variability be phylogenetic signal or stochastic noise). The resulting increase in likelihood is likely to dwarf the small penalty and result in an improved BIC score. We expect that if we generated a complementary cohort of alignments without the rate heterogeneity component, compared to ModelRevelator ML+BIC would have made more Type I errors, that is, falsely detecting the presence of rate heterogeneity in the data. We chose not to do this however, primarily because it is commonly recognised that rate homogeneous models are not expected to provide a good fit to empirical alignments. One can also envisage that the signal for rate heterogeneity would not have been strong in many alignments, due to the short sequence lengths in the PANDIT data, and possible high alpha values in some datasets (corresponding to weak rate heterogeneity).

With respect to the recovered model of sequence evolution (ignoring whether or not rate heterogeneity was recommended), ModelRevelator recovers the correct GTR-model for 75.49% of alignments, whereas ML+BIC recovers the correct model for 69.74% of alignments. However, the intersection of these groups, where both methods identified GTR, amounts to only 54.67% of the alignments. This means that the correct generative model was solely identified by ModelRevelator in 20.85% of alignments, and solely identified by ML+BIC in 15.08% of alignments. Interestingly, the two methods clearly diverge on the type of error they make when the alternate method is correct. When NNmodelfind is solely correct, ML+BIC predominantly selects K2P or HKY, and to a lesser extent TN93. These are all models which contain differential pairwise substitution rates between nucleotides. Conversely, when ML+BIC is solely correct ModelRevelator predominantly selects F81 or JC, models that assume the same pairwise substitution rates between all nucleotides.

With respect to topological inference, we again calculated the normalised Robinson-Foulds distances between the simulation tree, *T*SIM, and the trees inferred under the models selected by ModelRevelator and ML+BIC, 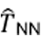 and *T*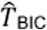 respectively. As before, where ModelRevelator recommended the inclusion of a rate heterogeneous model, the estimated alpha parameter was fixed during the inference, so that only other model parameters and edge lengths were optimised. In order to more clearly visualise any difference between the methods, we omitted the alignments for which both methods resulted in the same topology (64.4%). This left 22,990 (35.6%) alignments for which the two methods returned different trees. Figure 9 displays boxplots of normalised Robinson-Foulds distances for these alignments. The boxplots show that the results are similar between the two methods, although a marginal advantage to using ModelRevelator over ML+BIC is observed as sequence length increases. Given the fact that the PANDIT datasets were dissimilar to the Lanfear training data (in terms of distribution of generating parameters), this is a very encouraging sign that ModelRevelator may perform well on new empirical alignments generally.

**Figure 9:**
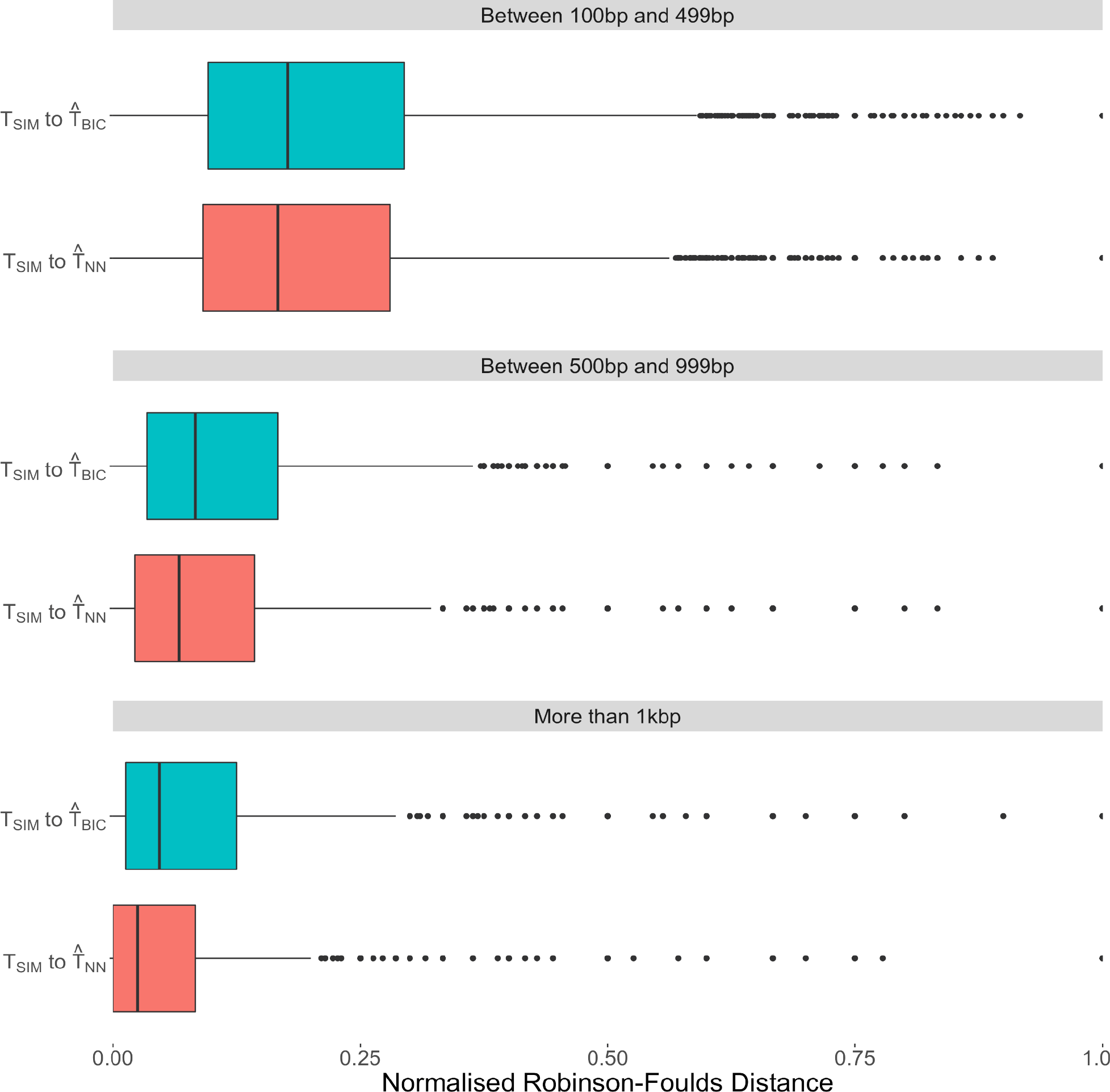
Robinson-Foulds distances to the simulation tree for the PANDIT-based simulations. Only MSAs for which use of ModelRevelator and ML+BIC led to discordant tree reconstructions were included. MSAs were binned into three groups according to sequence length. MSAs shorter than 100bp were omitted.

### Comparing ModelRevelator to ModelTeller

ModelTeller (Abadi et al. 2020) is another recent, machine learning based approach for model selection, based on random forests. We performed a comparison of ModelTeller and ModelRevelator using the Lanfear test data (for details on the test data, see Methods), to see how accurately each method was able to recover the model used to simulate an alignment. As ModelTeller supports 24 models of sequence evolution (JC, K2P/K80, F81, HKY, SYM, GTR, with modifications +I, +G, +I+G) and ModelRevelator supports 12 models of sequence evolution, for our comparison, we chose the intersection of what is supported by both methods (10 models of sequence evolution). When comparing ModelTeller estimates to the true model of sequence evolution of the simulated data, we found that ModelTeller rarely returned the correct model of sequence evolution (Fig. 10; Supplementary fig. 11-14). Furthermore, the confusion matrices show a strong dependency on the number of taxa in the MSAs (Supplementary fig. 11-14), as the number of taxa increases so does the complexity of the models recommended by ModelTeller.

**Figure 10:**
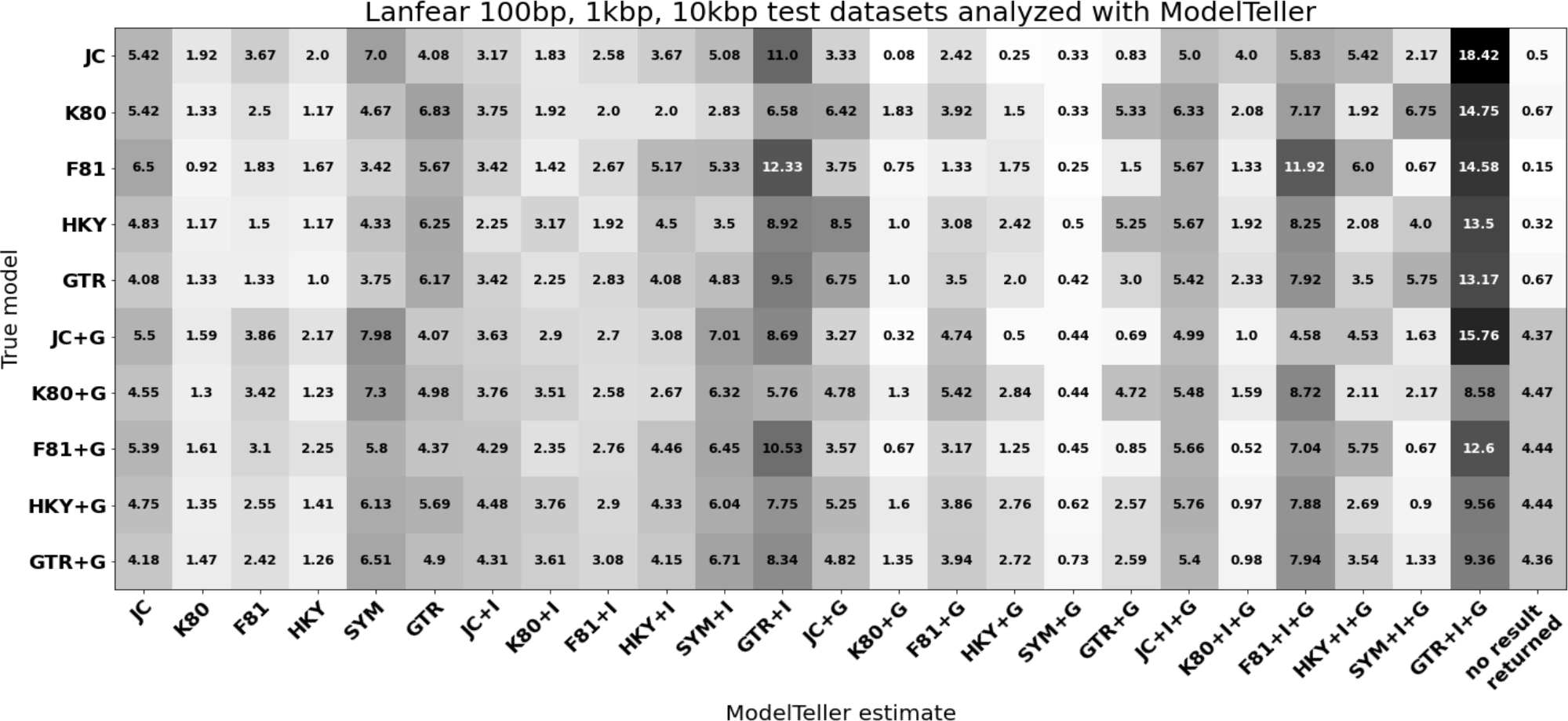
Results for ModelTeller on our Lanfear test data of 100bp, 1kbp and 10kbp MSAs. The rows show the true simulation model of the Lanfear test data and the columns show the percent of alignments that ModelTeller estimated each model. The final column displays the number of alignments for which ModelTeller analysis failed.

Figure 10 also shows that there is very little consistency in the models recommended by ModelTeller. In fact, the greatest success rate in correctly identifying the simulation model came for GTR-simulated alignments, at just 6.17%. No particular error patterns are obvious, rather the models chosen by ModelTeller appear to be somewhat random. It is important to make clear the distinction between the two methods at this point. ModelRevelator is trained to identify the model that best fits the data. On the other hand, ModelTeller aims to choose the model that leads to the most accurate branch length estimates. It makes no claim on being able to recover the model used to simulate the data. Therefore, its failure to reliably do so is not unexpected and should not be viewed as a criticism of the method’s performance. The two methods have different goals, and this should be kept in mind when making comparisons between them. With this in mind we further compared the two methods based on the accuracy of the trees (in terms of both topology and branch length estimates) that are inferred using the recommended models. For topological accuracy we compared the normalised Robinson-Foulds distances between the simulation tree (*T*_*SIM*_ and the trees inferred under models recommended by ModelTeller 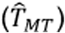 and ModelRevelator 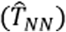. Similarly, to compare branch length estimates we calculated the branch score distance (Kuhner and Felsenstein 1994) instead of the normalised Robinson-Foulds distance. Supplementary Figures 15 and 16 indicate that both methods are similarly accurate with respect to both topology and branch length estimates.

These results indicate that while ModelTeller and ModelRevelator both lead to comparable tree reconstruction and branch length estimates (for these data at least), only ModelRevelator is able to infer the substitution model used to simulate the data to a reasonable degree of accuracy.

### Time measurements for ModelRevelator vs ML+BIC

Given a fixed input size, the computational expense of inference using a trained neural network is only dependent on the size of the network (number of parameters and operations).

Consequently, the time required for performing inference with NNmodelfind and NNalphafind is independent of alignment size. Considering ModelRevelator in its entirety however, the same cannot be said. Prior to carrying out inference with the networks, the MSA needs to be preprocessed in order to convert it into the correct format for input. As described in the methods section, this entails calculating summary statistics for randomly sampled taxon pairs, and the computation cost of this step grows with sequence length. By comparison, computation time for ML+BIC theoretically grows linearly with sequence length, and exponentially with number of taxa in the alignment. In Figure 11, we show the computation time for ModelRevelator (MSA preprocessing + NNmodelfind inference + NNalphafind inference) and the ML+BIC method.

**Figure 11:**
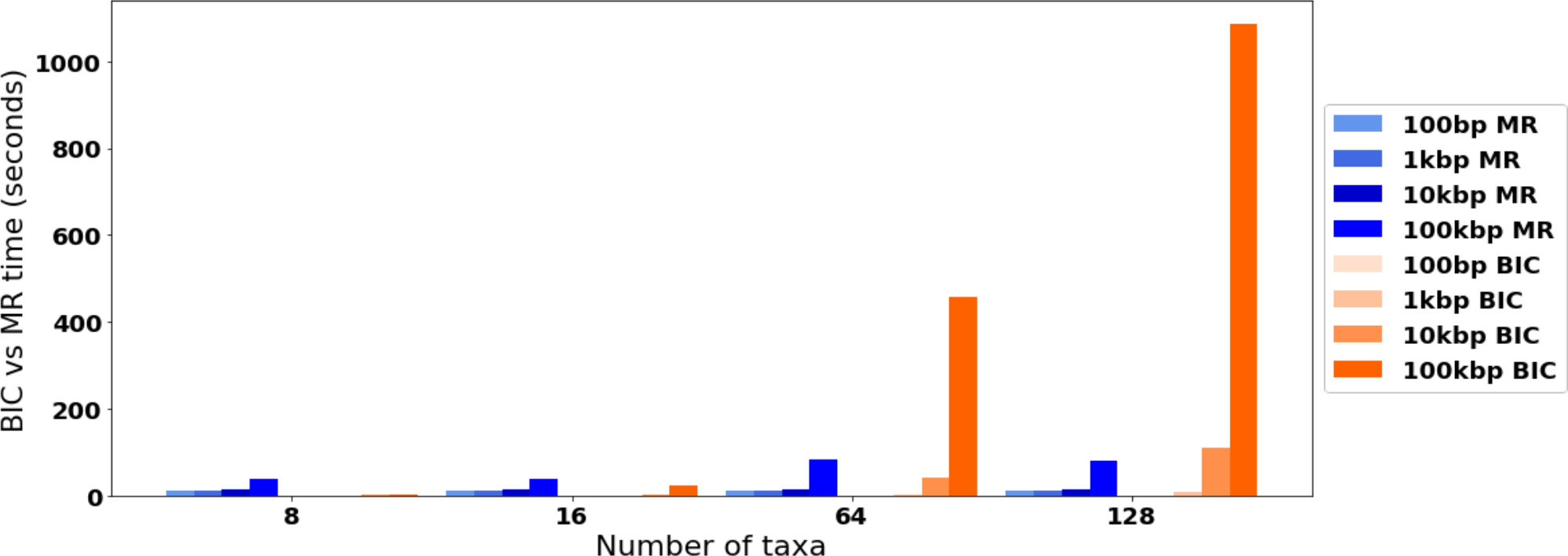
Mean computation time for ModelRevelator (MR) (MSA preprocessing + NNmodelfind + NNalphafind) vs ML+BIC (BIC) (full model estimation, including rate parameter estimation).

We observe that for small MSAs (100bp to 1kbp and 8 to 16 taxa), ML+BIC is typically faster than ModelRevelator, but the difference is somewhat negligible. For large alignments (100kbp and 128 taxa) ModelRevelator is approximately 14 times faster than ML+BIC. In a real world application setting, highly parallel neural network inference would be performed on a GPU or several CPU cores. Also, for partition models, neural network batch inference would allow for inference of many partitions at the same time.

## Discussion

The neural network-based ModelRevelator appears to perform comparably to the well-established approach of using ML inference and an information criterion (in this case BIC) to discriminate between models. Phylogenetic estimation was found to marginally improve using the neural networks compared to ML+BIC, with the additional benefit of significant potential savings in computation time, depending on the size of the alignment.

Encouragingly, both neural networks which underpin ModelRevelator were found to generalise well when tested on alignments that differed to those on which they were trained. No appreciable deterioration in performance was observed when performing estimation on longer sequences, or with larger numbers of taxa. Although we observe that, compared to ML+BIC, NNmodelfind does not extract information as well from longer alignments, this does not have an impact on tree topology. Additionally, our method outperformed ML+BIC (in terms of accuracy of phylogenetic reconstruction) when tested on the PANDIT alignments, a set of empirical alignments that the networks had not seen during training. This finding is of particular interest, as it is an indication that ModelRevelator can be used with confidence by the community more broadly.

One area in which we envisage a particular benefit of our approach is in conjunction with partition models. Large alignments are often partitioned by gene, and/or codon position, into hundreds or even thousands of independent blocks. PartitionFinder (Lanfear et al. 2012) relies on information criteria to merge blocks that can be effectively modelled together, and selects models of sequence evolution to be applied to each block. Selecting a model on a large number of blocks can be accomplished efficiently by NNmodelfind running on a GPU, but ML+BIC would require parallelisation on available CPU cores. Depending on the hardware available, it is likely that the ML+BIC would be significantly slower for large projects.

Before deciding on the final architecture for NNmodelfind and NNalphafind, we experimented widely with a range of architectures. For NNmodelfind we tried conventional multi-layer perceptrons and convolutional neural networks as architectures, before settling on the final ResNet-18 architecture (He et al. 2016). For NNalphafind, we tried LSTMs with or without convolution layers for encoding, or with or without the attention layer. We found that both the convolution encoding layers and the attention layer were crucial for achieving reasonable estimates for alpha. We also explored using the ResNet-18 to estimate alpha but the results were not satisfactory. Another aspect which lent itself to experimentation was how to represent MSAs of varying size and length so that they could be input to the networks. We restricted ourselves to a fixed input size for both networks, but experimented with different input sizes, as well as different ways of transforming variably-sized MSAs into a fixed size input. An obvious drawback of the fixed size input is that it is not always possible for the network to utilise all the information available in large alignments. A promising avenue to address this might be networks that accept graph-based representations of MSAs as their input, as explored in (Drucks 2021).

The only other tool currently employing a machine learning approach for model estimation is ModelTeller (Abadi et al. 2020). Our simulation-based analysis of ModelTeller suggests that it struggles to recover the actual model of sequence evolution, although that is not the primary aim of the software. When performing a direct comparison to ModelRevelator, we find that although the actual model of sequence evolution is rarely found by ModelTeller, there is no appreciable difference in the accuracy of the resulting tree reconstructions. However, the focus of both methods is different. ModelRelevator is concerned with estimating the model of sequence evolution and the shape parameter alpha of the Γ-distribution, and is trained and tested only on this basis. ModelTeller focuses on finding the model that yields the most accurate edge lengths, and is trained and tested on this basis.

While some have argued that model selection is not a critical step in phylogenetic reconstruction (Abadi et al. 2020), this view is by no means the consensus (Ripplinger and Sullivan 2008; Hoff et al. 2016). It may be true that phylogenetic inference is robust to misspecified substitution models in a majority of cases, nevertheless it is in the minority of cases where this is not the case that model selection can be pivotal. Where there are contentious topologies with conflicting evidence, having confidence in a model selection methodology can add support to a hypothesis.

Notwithstanding our extensive explorations, we recognise the available options for potential neural network architectures is vast. It is therefore likely that neural net architectures exist which might yield better performance than those we present here. This fact, combined with the recognised shortcomings of information theoretic approaches, and the absence of directly comparable tools currently available, suggests that machine learning approaches to model selection represent fertile ground for ongoing investigation and development.

## Acknowledgements

We would like to especially thank Rob Lanfear for providing the empirical MSAs used in this study. AvH is supported by a grant from the FWF (FWF I 4686 – B).

## Notes

https://github.com/cibiv/ModelRevelator/

### Competing Interest Statement

The authors have declared no competing interest.

### Summary of Updates

We included extensive comparison to ModelTeller, a machine learning framework also intended for phylogenetic model estimation. Also, we performed comprehensive analysis of the quality of the phylogenetic trees produced, based on the phylogenetic models suggested by each method. For this, we evaluated tree topologies and branch lengths resulting from using the suggested models of sequence evolution.

