## supplementary material for "ModelRevelator: Fast phylogenetic model estimation via deep learning"

#### Supplementary Figures

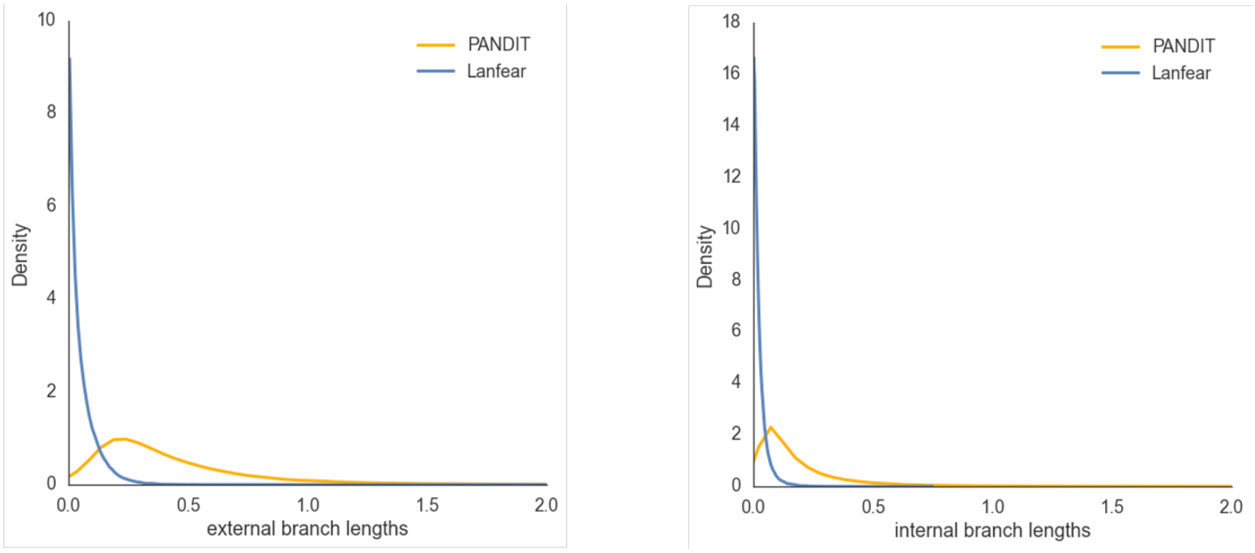


Supplementary Figure 1: Distribution of external and internal edge lengths of Lanfear and PANDIT data.

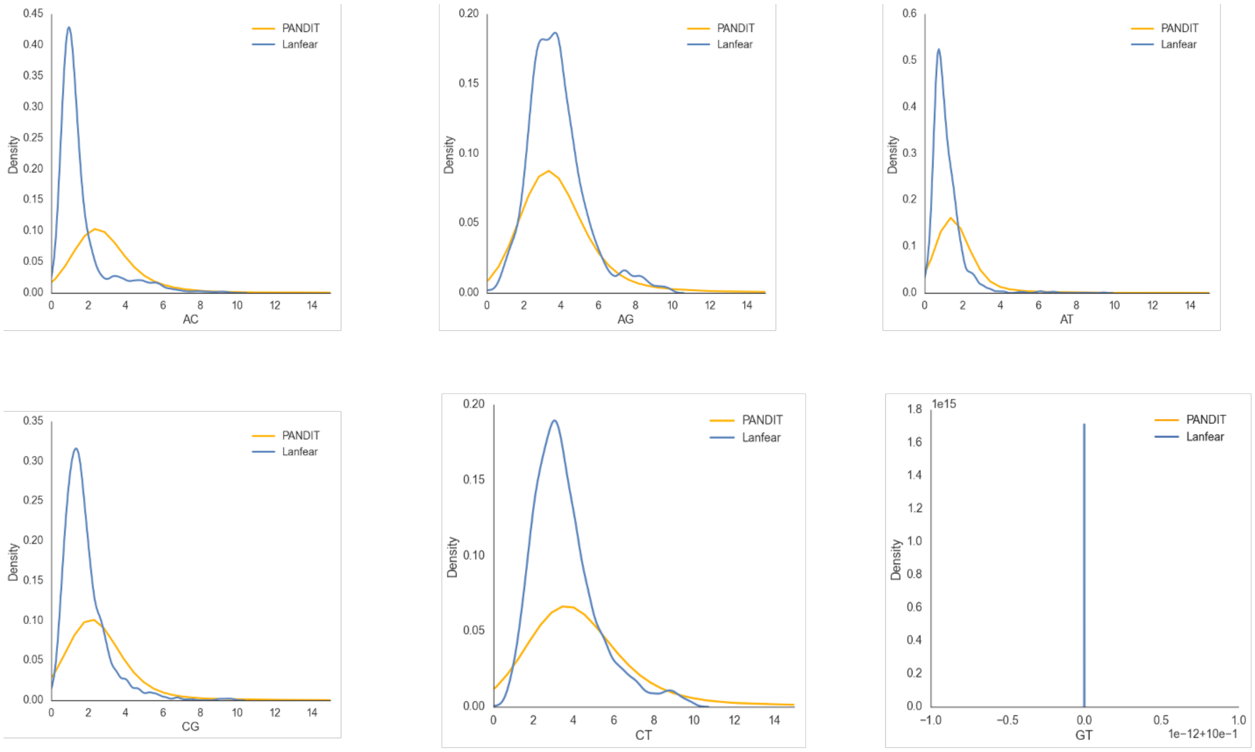

Supplementary Figure 2: Distributions of the 6 substitution rates for the two empirical datasets Lanfear and PANDIT. Smoothing of the histograms was achieved using kernel density estimation. As it is common practice, the relative substitution rates are normalised such that the GT rate is 1 in all datasets.

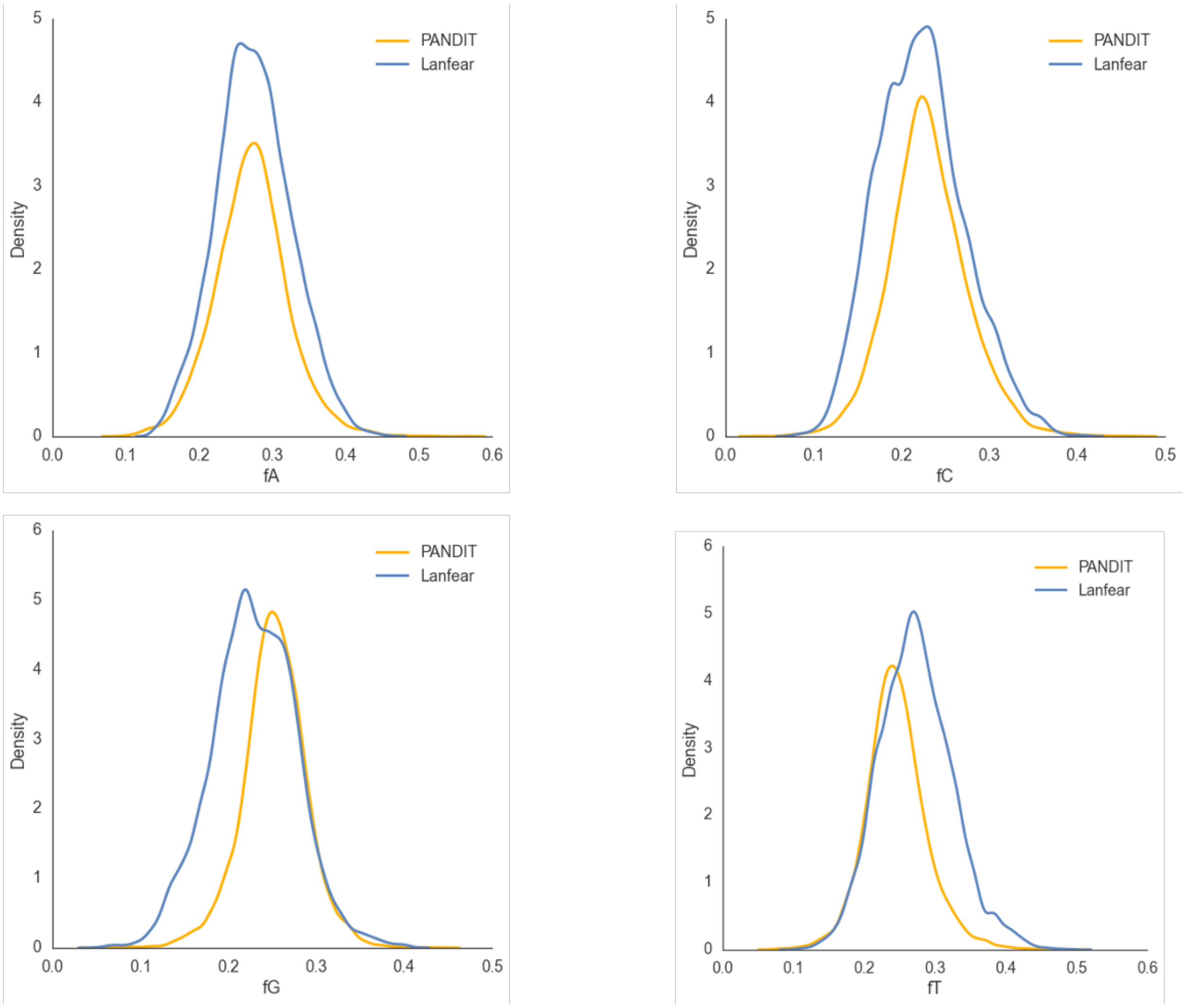


Supplementary Figure 3: Distributions of the 4 base frequencies for the two empirical datasets Lanfear and PANDIT. Smoothing of the histograms was achieved using kernel density estimation.

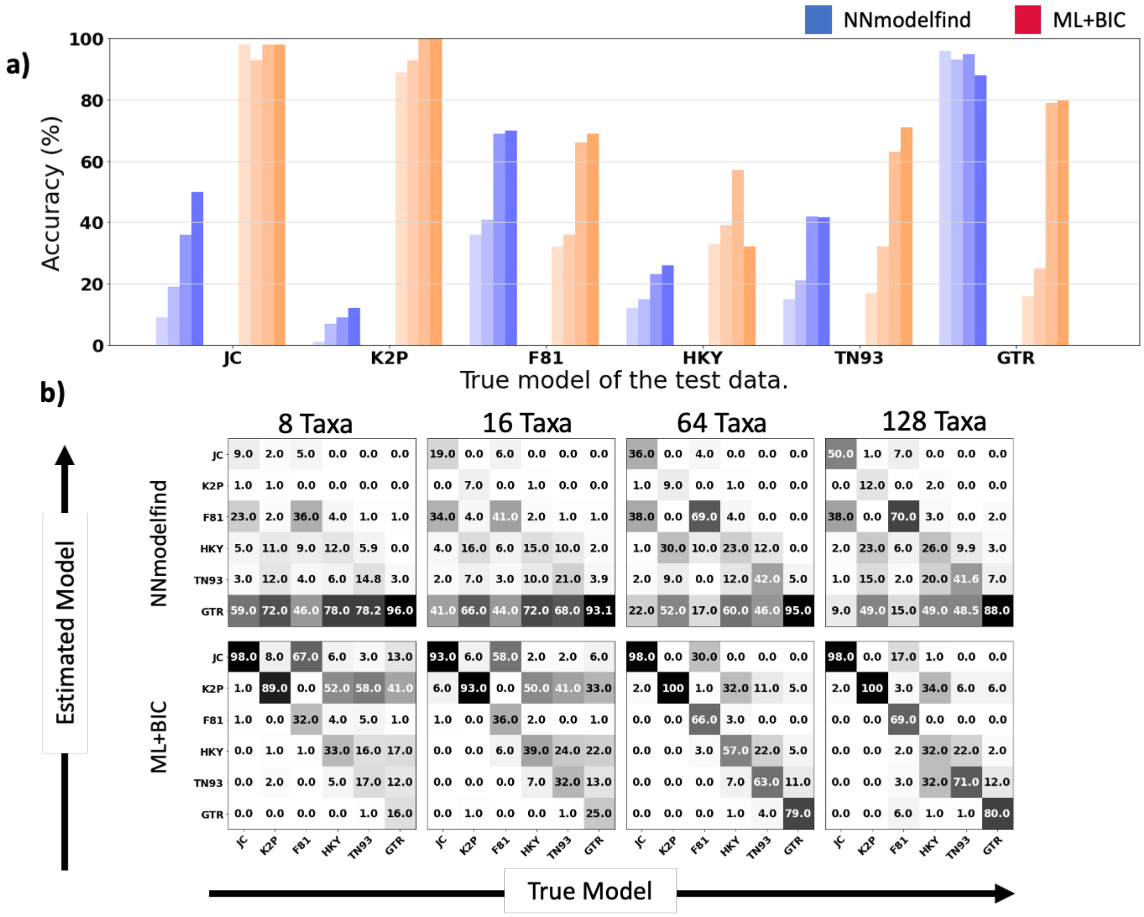


Supplementary Figure 4: Model selection accuracy of NNmodelfind and ML+BIC on 100bp long Rhom MSAs. **(A)** Blueish colours indicate results from NNmodelfind, reddish colours indicate results from ML+BIC. Darkening bars distinguish increasing number of taxa from left to right (8, 16, 64 and 128). **(B)** Confusion matrices for NNmodelfind and ML+BIC. Entries along the diagonal indicate the percentage of alignments for which the correct model was identified.

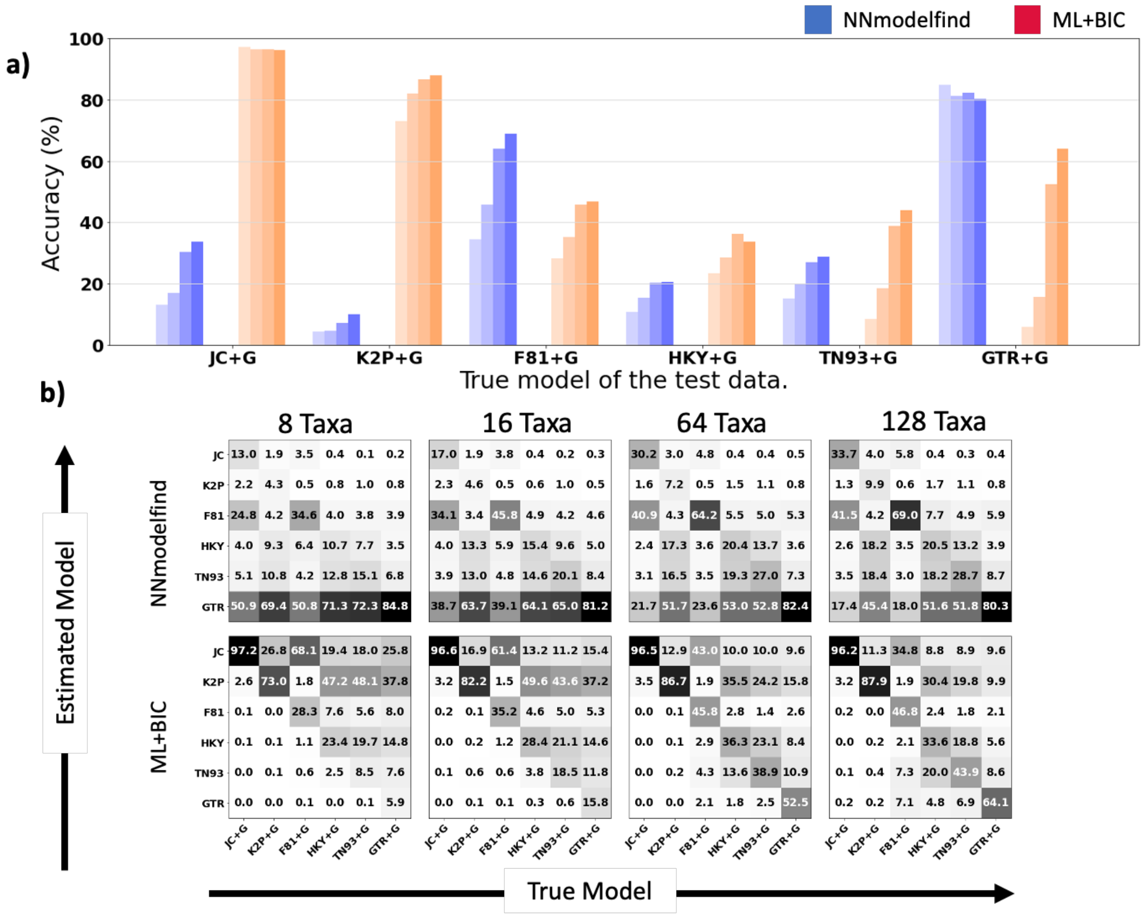


Supplementary Figure 5: Model selection accuracy of NNmodelfind and ML+BIC on 100bp long Rhet MSAs. **(A)** Blueish colours indicate results from NNmodelfind, reddish colours indicate results from ML+BIC. Darkening bars distinguish increasing number of taxa from left to right (8, 16, 64 and 128). **(B)** Confusion matrices for NNmodelfind and ML+BIC. Entries along the diagonal indicate the percentage of alignments for which the correct model was identified.

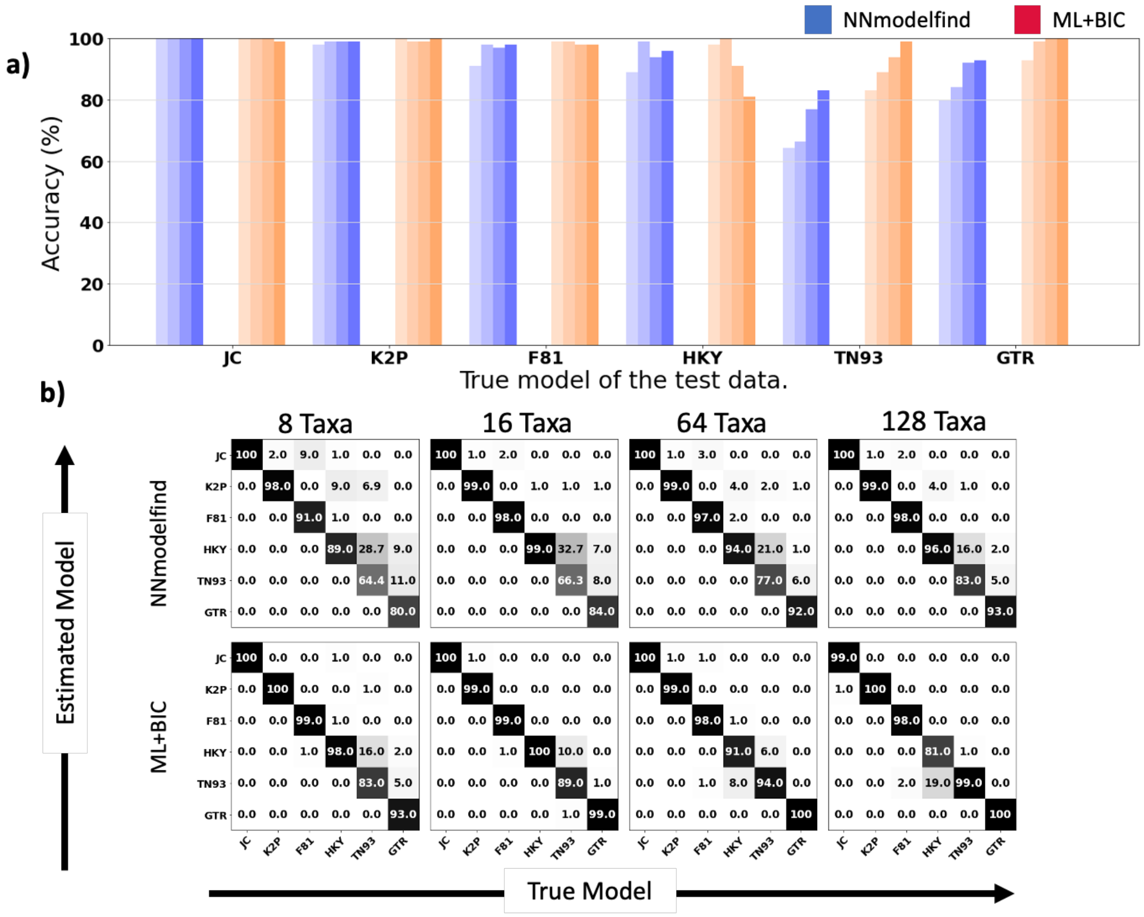


Supplementary Figure 6: Model selection accuracy of NNmodelfind and ML+BIC on 10kbp long Rhom MSAs. **(A)** Blueish colours indicate results from NNmodelfind, reddish colours indicate results from ML+BIC. Darkening bars distinguish increasing number of taxa from left to right (8, 16, 64 and 128). **(B)** Confusion matrices for NNmodelfind and ML+BIC. Entries along the diagonal indicate the percentage of alignments for which the correct model was identified.

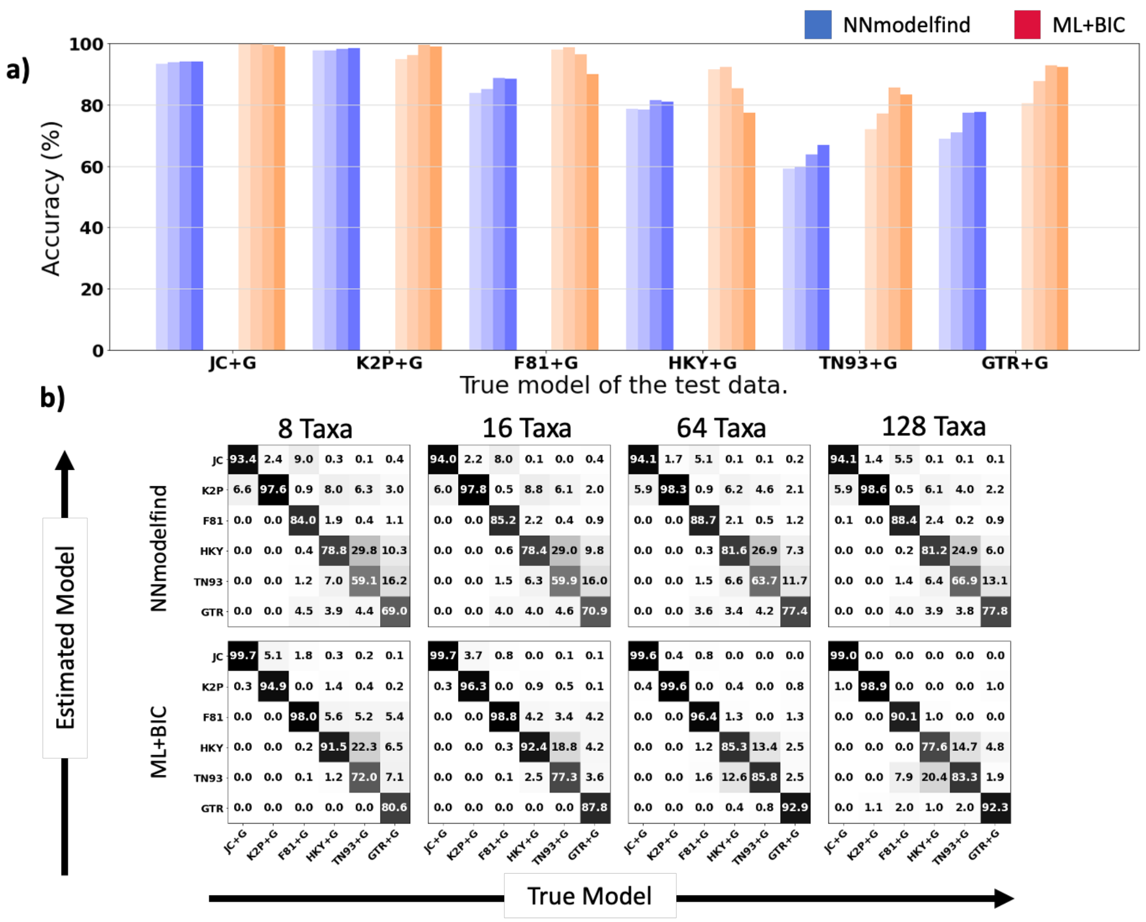


Supplementary Figure 7: Model selection accuracy of NNmodelfind and ML+BIC on 10kbp long, Rhet MSAs. **(A)** Blueish colours indicate results from NNmodelfind, reddish colours indicate results from ML+BIC. Darkening bars distinguish increasing number of taxa from left to right (8, 16, 64 and 128). **(B)** Confusion matrices for NNmodelfind and ML+BIC. Entries along the diagonal indicate the percentage of alignments for which the correct model was identified.

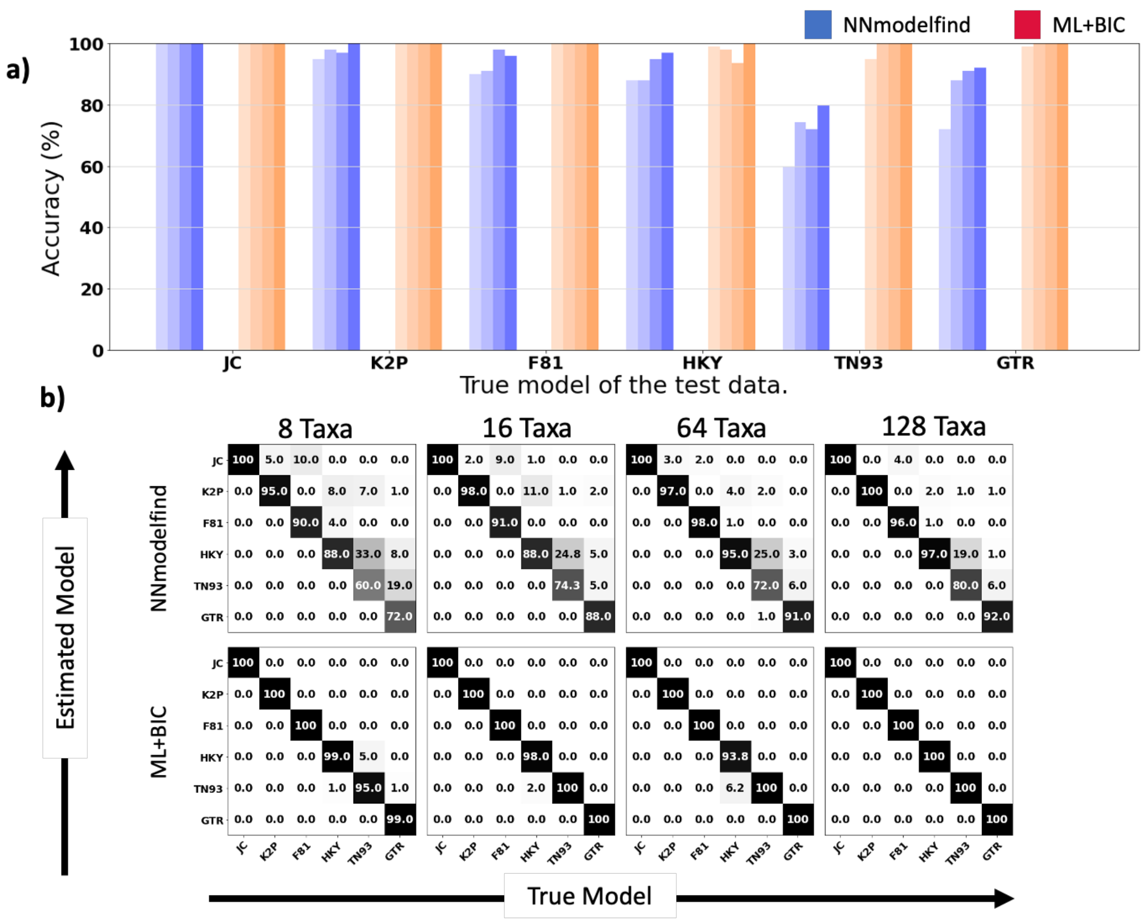


Supplementary Figure 8: Model selection accuracy of NNmodelfind and ML+BIC on 100kbp long Rhom MSAs. **(A)** Blueish colours indicate results from NNmodelfind, reddish colours indicate results from ML+BIC. Darkening bars distinguish increasing number of taxa from left to right (8, 16, 64 and 128). **(B)** Confusion matrices for NNmodelfind and ML+BIC. Entries along the diagonal indicate the percentage of alignments for which the correct model was identified.

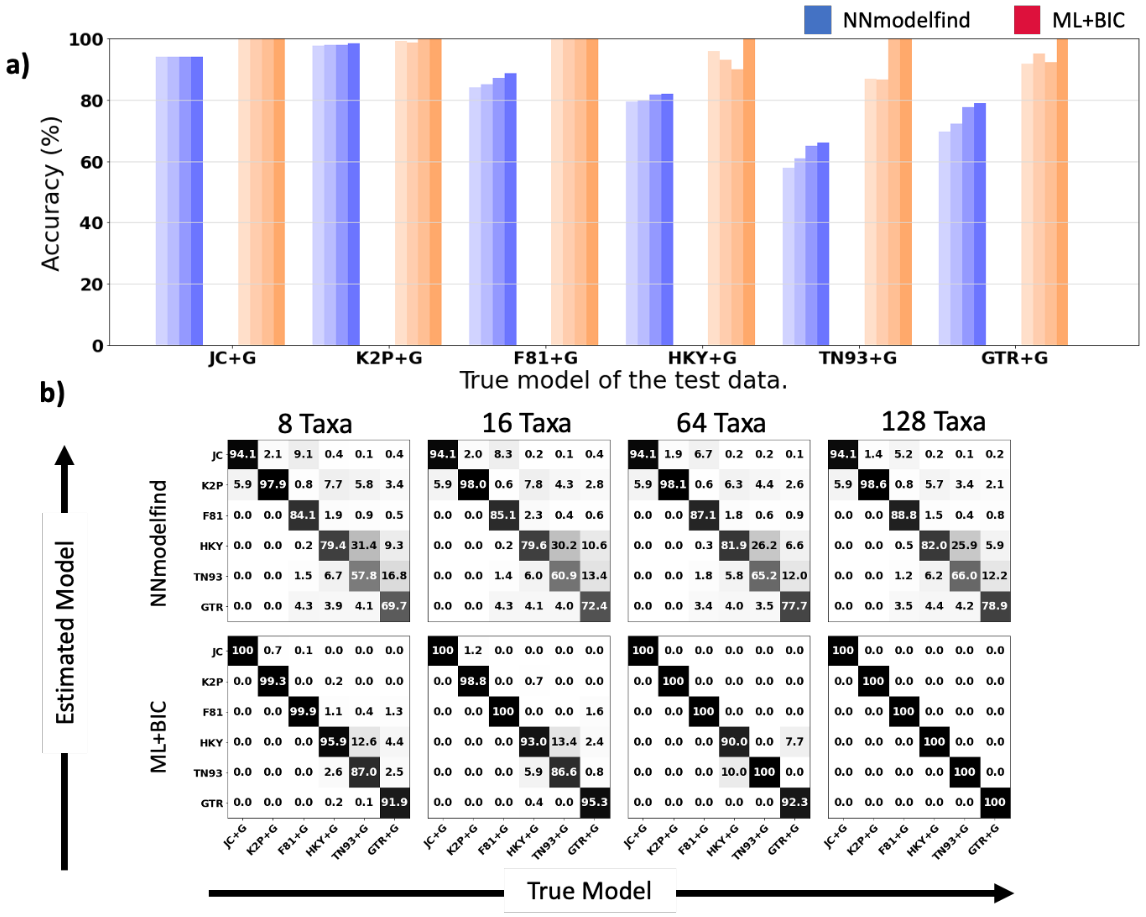


Supplementary Figure 9: Model selection accuracy of NNmodelfind and ML+BIC on 100kbp long Rhet MSAs. **(A)** Blueish colours indicate results from NNmodelfind, reddish colours indicate results from ML+BIC. Darkening bars distinguish increasing number of taxa from left to right (8, 16, 64 and 128). **(B)** Confusion matrices for NNmodelfind and ML+BIC. Entries along the diagonal indicate the percentage of alignments for which the correct model was identified.

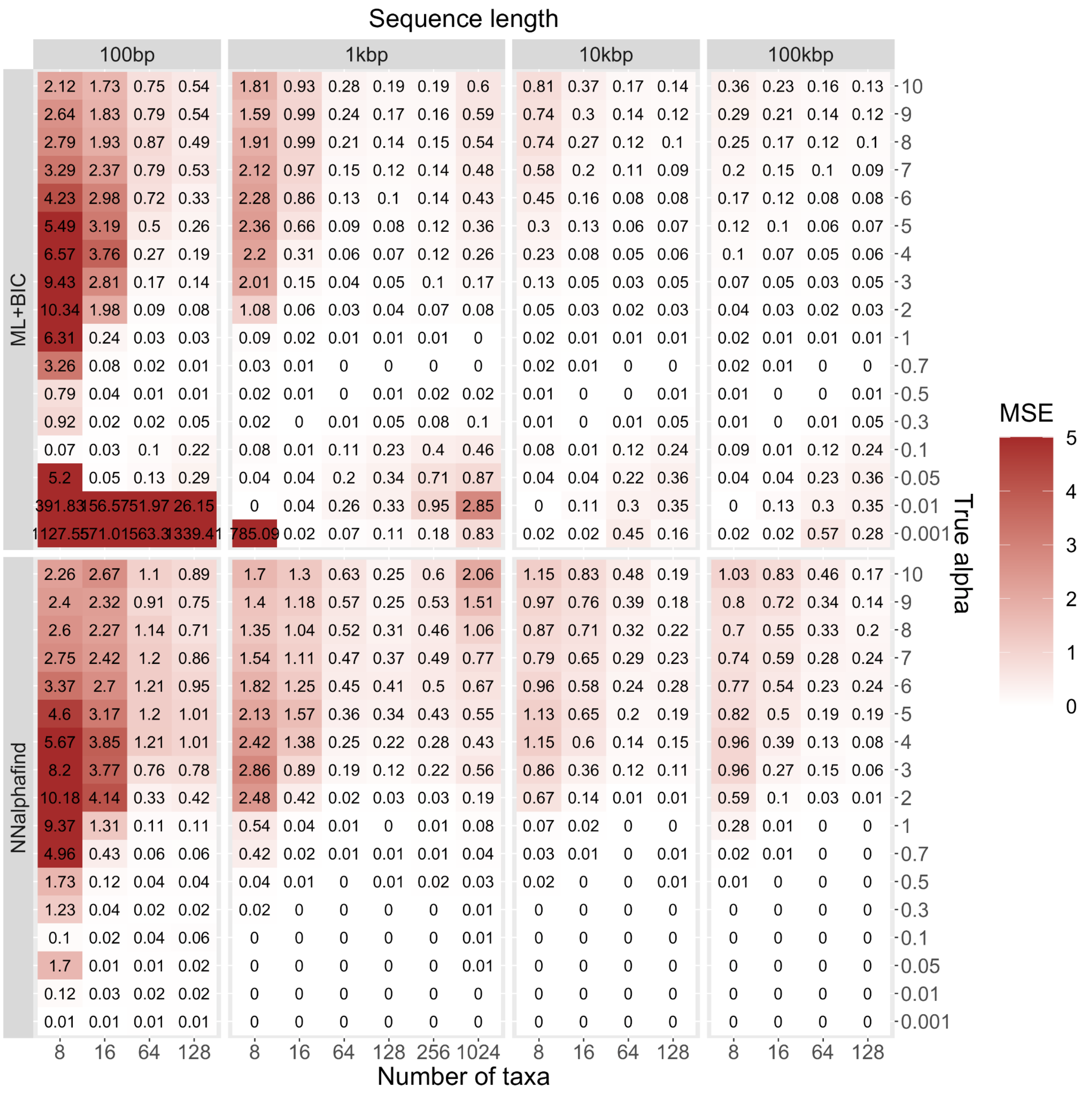


*Supplementary Figure 10: Mean squared error of inferred alpha value and true alpha value, stratified by inference method (ML+BIC or NNalphafind), number of taxa (8, 16, 64, and 128), sequence length (100bp, 1kbp, 10kbp, and 100kbp), and true alpha value (17 levels ranging from 0.001 (very strong heterogeneity) up to 10 (very weak heterogeneity)).*

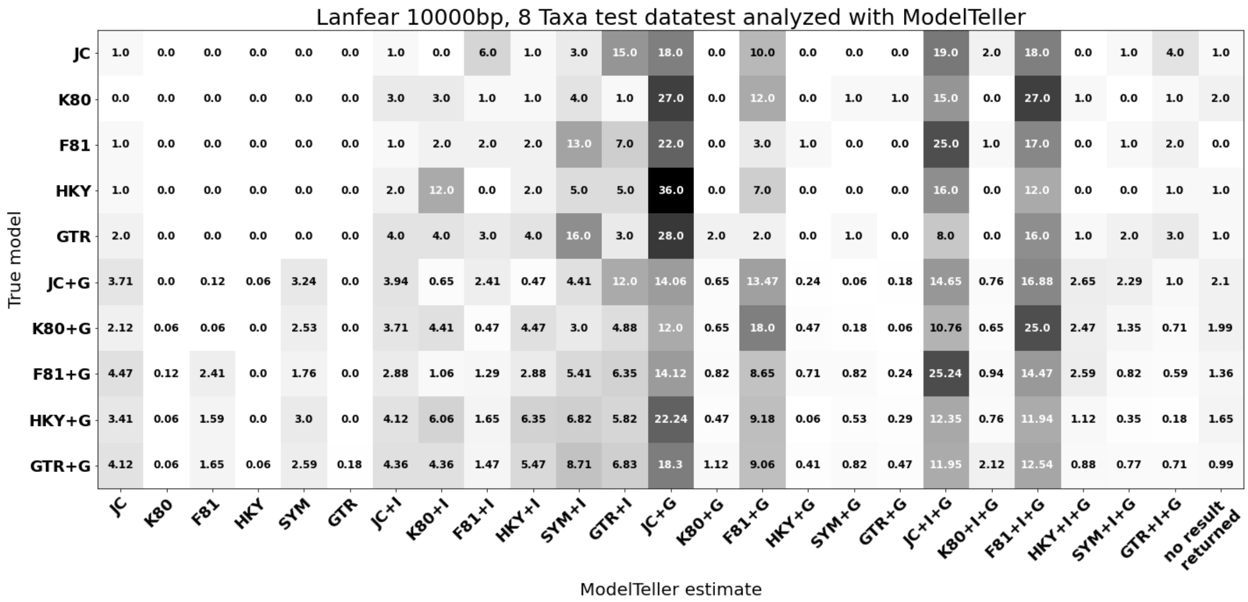


Supplementary Figure 11: Results for ModelTeller on our Lanfear test data for 8 taxa, 10kbp MSAs. The rows show the true simulation model of the Lanfear test data and the columns show the percent of alignments that ModelTeller estimated for each model. The final column displays the number of alignments for which ModelTeller analysis failed.

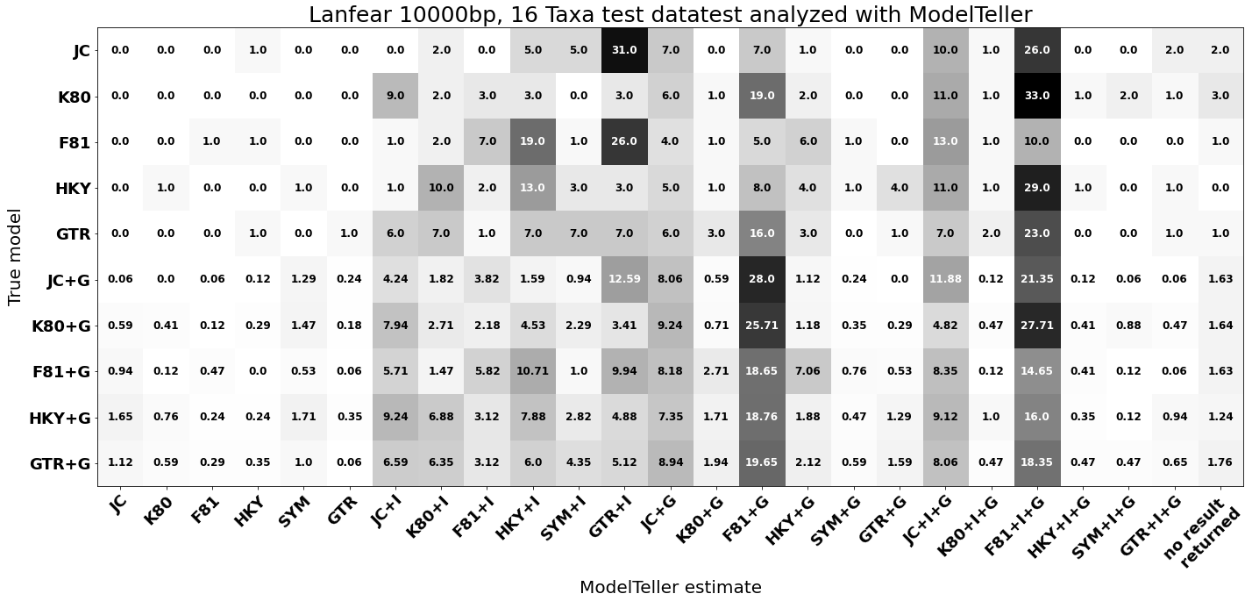


Supplementary Figure 12: Results for ModelTeller on our Lanfear test data for 16 taxa, 10kbp MSAs. The rows show the true simulation model of the Lanfear test data and the columns show the percent of alignments that ModelTeller estimated for each model. The final column displays the number of alignments for which ModelTeller analysis failed.

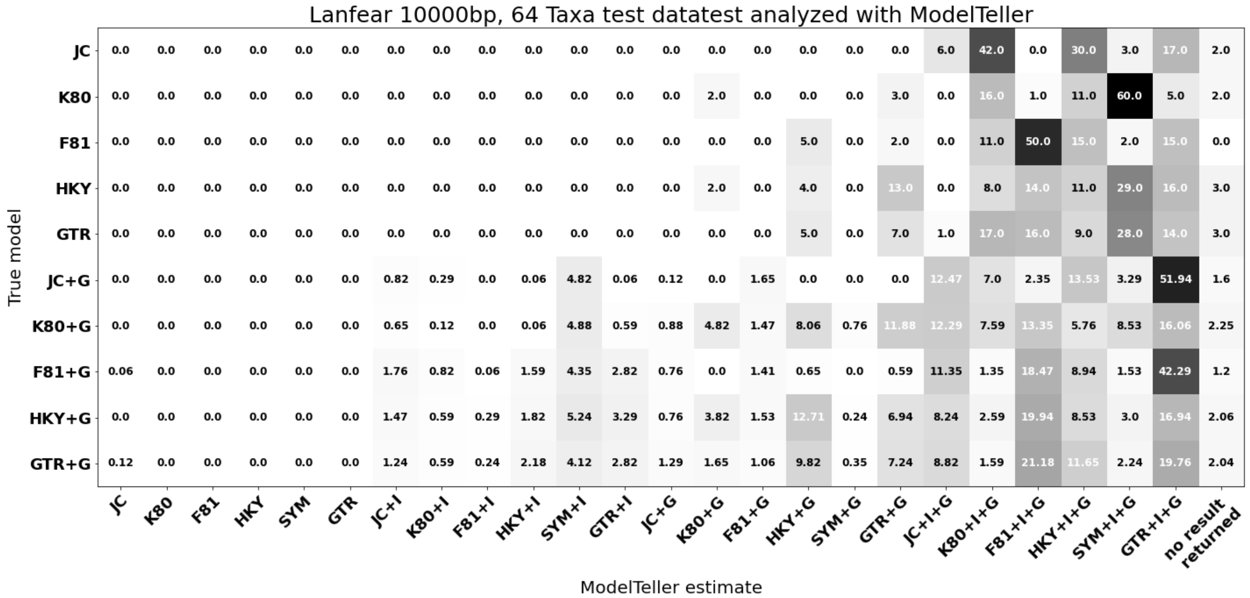


Supplementary Figure 13: Results for ModelTeller on our Lanfear test data for 64 taxa, 10kbp MSAs. The rows show the true simulation model of the Lanfear test data and the columns show the percent of alignments that ModelTeller estimated for each model. The final column displays the number of alignments for which ModelTeller analysis failed.

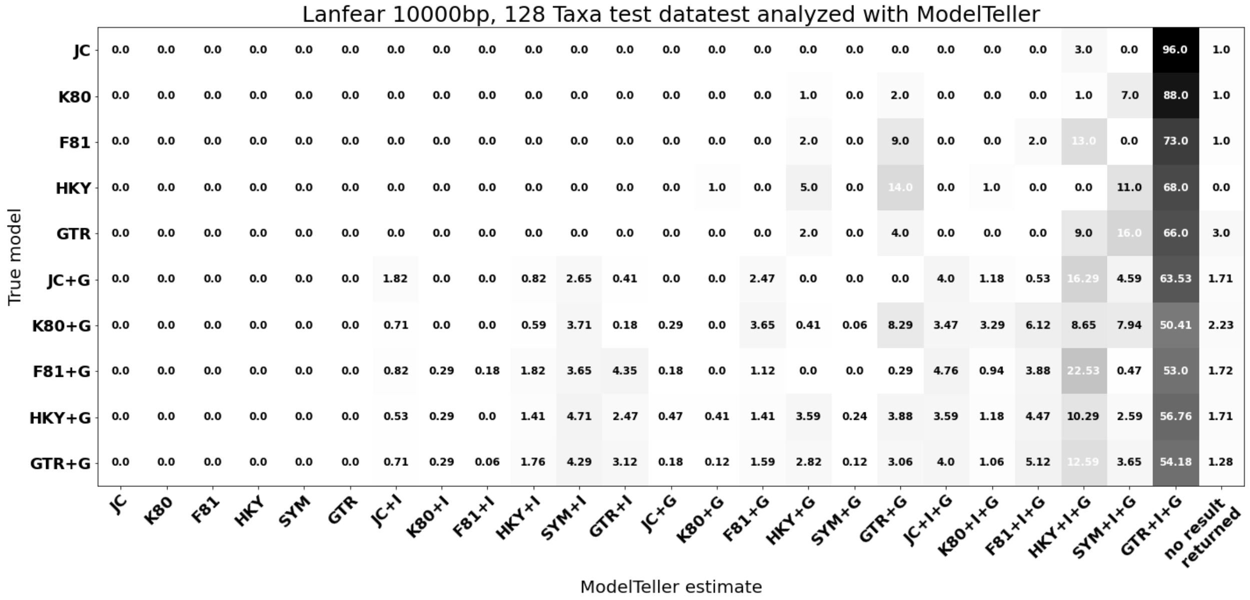

Supplementary Figure 14: Results for ModelTeller on our Lanfear test data for 128 taxa, 10kbp MSAs. The rows show the true simulation model of the Lanfear test data and the columns show the percent of alignments that ModelTeller estimated for each model. The final column displays the number of alignments for which ModelTeller analysis failed.

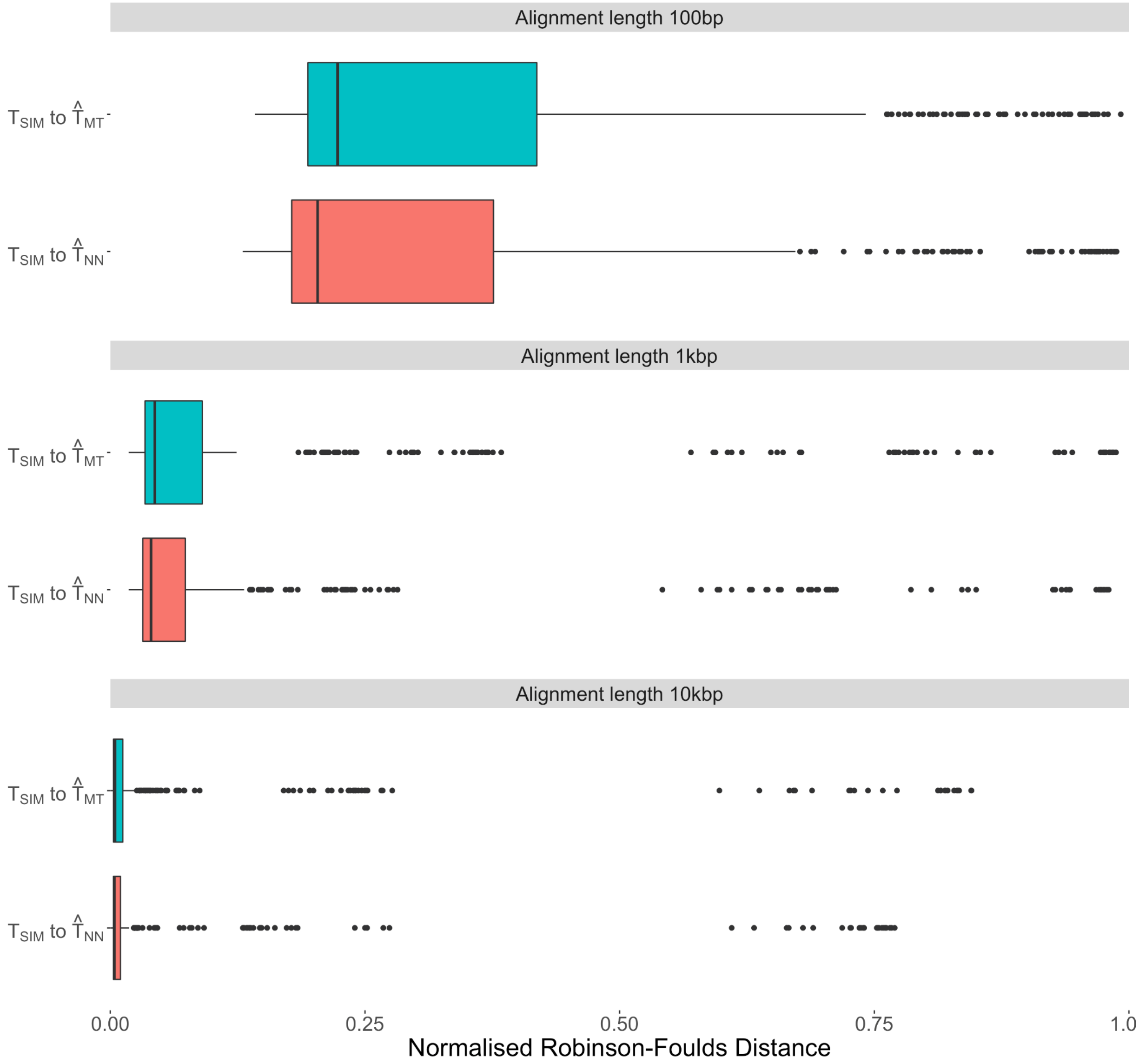


Supplementary Figure 15: Boxplots of mean normalised Robinson-Foulds distance between the simulation tree and the trees inferred based on model selection via ModelRevelator (NN, red)

and ModelTeller (MT, blue). The dataset used for comparison was the Lanfear-based simulated test data.

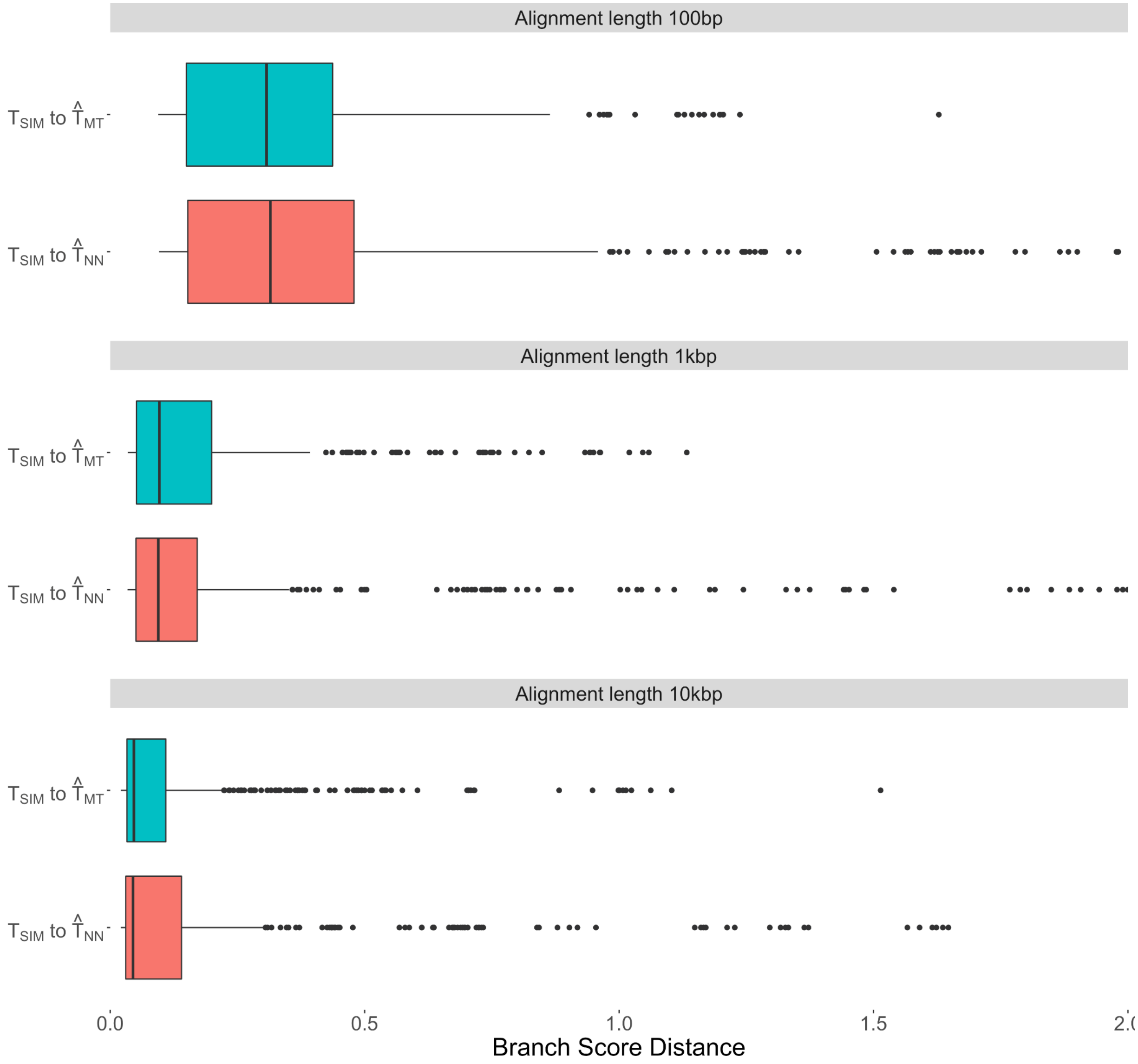


Supplementary Figure 16: Boxplots of mean branch score distance between the simulation tree and the trees inferred based on model selection via ModelRevelator (NN, red) and ModelTeller (MT, blue). The dataset used for comparison was the Lanfear-based simulated test data.

### Source information for Lanfear alignments

Alignments extracted from the Lanfear collection which formed the basis of the simulated training and test data originated in the following 31 publications.


1. Anderson FE, Bergman A, Cheng SH, Pankey MS, Valinassab T. Lights out: the evolution of bacterial bioluminescence in Loliginidae. Hydrobiologia. 2014 Mar 1;725(1):189-203.
2. Bergsten J, Nilsson AN, Ronquist F. Bayesian tests of topology hypotheses with an example from diving beetles. Systematic biology. 2013 Sep 1;62(5):660-73.
3. Branstetter MG, Danforth BN, Pitts JP, Faircloth BC, Ward PS, Buffington ML, Gates MW, Kula RR, Brady SG (2017) Phylogenomic Insights into the Evolution of Stinging Wasps and the Origins of Ants and Bees. Current Biology 27(7): 1019-1025.
4. Broughton RE, Betancur RR, Li C, Arratia G, Ortí G. Multi-locus phylogenetic analysis reveals the pattern and tempo of bony fish evolution. PLoS Curr. 2013 Apr 16;5(5).
5. Brown RM, Siler CD, Das I, Min Y. Testing the phylogenetic affinities of Southeast Asia’s rarest geckos: Flap-legged geckos (Luperosaurus), Flying geckos (Ptychozoon) and their relationship to the pan-Asian genus Gekko. Molecular phylogenetics and evolution. 2012 Jun 30;63(3):915-21.
6. Cannon JT, Vellutini BC, Smith J, Ronquist F, Jondelius U, Hejnol A (2016) Xenacoelomorpha is the sister group to Nephrozoa. Nature 530(7588): 89–93. http://dx.doi.org/10.1038/nature16520
7. Cognato AI, Vogler AP. Exploring data interaction and nucleotide alignment in a multiple gene analysis of Ips (Coleoptera: Scolytinae). Systematic Biology. 2001 Nov 1;50(6):758-80.
8. Day JJ, Peart CR, Brown KJ, Friel JP, Bills R, Moritz T. Continental diversification of an African catfish radiation (Mochokidae: Synodontis). Systematic biology. 2013 Jan 9:syt001.
9. Devitt TJ, Devitt SE, Hollingsworth BD, McGuire JA, Moritz C. Montane refugia predict population genetic structure in the Large-blotched Ensatina salamander. Molecular ecology. 2013 Mar 1;22(6):1650-65.
10. Dornburg A, Moore JA, Webster R, Warren DL, Brandley MC, Iglesias TL, Wainwright PC, Near TJ. Molecular phylogenetics of squirrelfishes and soldierfishes (Teleostei:

Beryciformes: Holocentridae): Reconciling more than 100 years of taxonomic confusion. Molecular phylogenetics and Evolution. 2012 Nov 30;65(2):727-38.

11. Faircloth BC, Sorenson L, Santini F, Alfaro ME (2013) A phylogenomic perspective on the radiation of ray-finned fishes based upon targeted sequencing of ultraconserved elements (UCEs). PLoS ONE 8(6): e65923
12. Fong JJ, Brown JM, Fujita MK, Boussau B. A phylogenomic approach to vertebrate phylogeny supports a turtle-archosaur affinity and a possible paraphyletic lissamphibia. PLoS One. 2012 Nov 7;7(11):e48990.
13. Horn JW, Xi Z, Riina R, Peirson JA, Yang Y, Dorsey BL, Berry PE, Davis CC, Wurdack KJ (2014) Evolutionary bursts in Euphorbia (Euphorbiaceae) are linked with photosynthetic pathway. Evolution 68(12): 3485-3504.
14. Kawahara AY, Rubinoff D. Convergent evolution of morphology and habitat use in the explosive Hawaiian fancy case caterpillar radiation. Journal of evolutionary biology. 2013 Aug 1;26(8):1763-73.
15. Lartillot N, Delsuc F. Joint reconstruction of divergence times and life-history evolution in placental mammals using a phylogenetic covariance model. Evolution. 2012 Jun 1;66(6):1773-87.
16. Looney BP, Ryberg M, Hampe F, Sánchez-García M, Matheny PB (2016) Into and out of the tropics: global diversification patterns in a hyper-diverse clade of ectomycorrhizal fungi. Molecular Ecology 25(2): 630–647.
17. Moyle RG, Oliveros CH, Andersen MJ, Hosner PA, Benz BW, Manthey JD, Travers SL, Brown RM, Faircloth BC. Tectonic collision and uplift of Wallacea triggered the global songbird radiation. Nature Communications. 2016 Aug 30;7.
18. Murray EA, Carmichael AE, Heraty JM. Ancient host shifts followed by host conservatism in a group of ant parasitoids. Proceedings of the Royal Society of London B: Biological Sciences. 2013 May 22;280(1759):20130495.
19. Near TJ, Dornburg A, Eytan RI, Keck BP, Smith WL, Kuhn KL, Moore JA, Price SA, Burbrink FT, Friedman M, Wainwright PC. Phylogeny and tempo of diversification in the superradiation of spiny-rayed fishes. Proceedings of the National Academy of Sciences. 2013 Jul 30;110(31):12738-43.
20. Oaks JR. A time-calibrated species tree of Crocodylia reveals a recent radiation of the true crocodiles. Evolution. 2011 Nov 1;65(11):3285-97.
